# Rala and the exocyst control Pvr trafficking and signaling to ensure lymph gland homeostasis in *Drosophila melanogaster*

**DOI:** 10.1101/2020.05.26.086140

**Authors:** Helene Knævelsrud, Caroline Baril, Gwenaëlle Gavory, Jorrit M. Enserink, Marc Therrien

## Abstract

The balance between hematopoietic progenitors and differentiated hemocytes is finely tuned during development. In the larval hematopoietic organ of *Drosophila*, called the lymph gland, the receptor tyrosine kinase Pvr signals from differentiated cells to maintain a pool of undifferentiated progenitors. However, little is known about the processes that support Pvr function. The small GTPase Ral is involved in the regulation of several membrane trafficking events. *Drosophila* has a single Ral protein, Rala, which has been implicated in the development of various tissues. Here, we investigated the involvement of Rala in the larval fly hematopoietic system. We discovered that the loss of *Rala* activity phenocopies *Pvr* loss of function by promoting hemocyte progenitor differentiation. Moreover, using epistasis analysis, we found that the guanine exchange factor RalGPS lies upstream of Rala in this event, whereas the exocyst and Rab11 are acting downstream. Strikingly, the loss of Rala activity leads to a considerable accumulation of Pvr at the plasma membrane, hence suggesting a trafficking defect and reduced Pvr function. Consistent with this hypothesis, *Rala* loss of function phenotype in the lymph gland is fully suppressed by constitutive STAT activity, which normally mediates Pvr function in the lymph gland. Together, our findings unravel a novel RalGPS-Rala-exocyst-Rab11 axis for the maintenance of lymph gland homeostasis through Pvr.

## Introduction

Maintaining an appropriate pool of hematopoietic progenitors and avoiding aberrant blood cell development is crucial for the fitness of an organism. The fruit fly *Drosophila* melanogaster is a genetically tractable model for signaling mechanisms controlling hematopoiesis (Banerjee, Girard, Goins, & Spratford, 2019). The evolutionary conservation of several of the signaling pathways and transcription factors underlying the development of its hematopoietic system also makes this model biomedically relevant (Boulet, Miller, Vandel, & Waltzer, 2018).

*Drosophila* hematopoiesis occurs in embryo and larvae, whereas the existence of hematopoiesis in adult flies is debated (Letourneau et al., 2016; Sanchez Bosch et al., 2019). A first wave of hematopoiesis starts with the specification of a group of cells derived from the head mesoderm anlage that will give rise to circulating hemocytes in the embryo. Later, the cells giving rise to the larval hematopoietic organ, the lymph gland, are specified from the cardiogenic mesoderm. The lymph gland consists of a pair of primary lobes and several secondary lobes aligned along the dorsal vessel. The primary lobe is subdivided into three compartments: the medullary zone (MZ) primarily composed of progenitor cells, the cortical zone (CZ) containing differentiated hemocytes and the posterior signaling centre (PSC), which regulates the balance between the two other compartments. Differentiated hemocytes (Honti, Csordas, Kurucz, Markus, & Ando, 2014) are mostly plasmatocytes, which phagocytose dying cells and invading pathogens. There is a small population of crystal cells, which perform melanization and are involved in wound healing. Specific challenges such as eggs of parasitic wasps will induce the differentiation of lamellocytes, which are large cells that work together with crystal cells to encapsulate their targets. The larval lymph gland is used as a model system to study hematopoietic progenitor balance (Banerjee et al., 2019). From the late second instar, the PSC secretes both Hh and Pvf1 as signals to maintain progenitor quiescence (Baldeosingh, Gao, Wu, & Fossett, 2018; Mandal, Martinez-Agosto, Evans, Hartenstein, & Banerjee, 2007; Mondal et al., 2011). The Hh signal acts on the MZ (Mandal et al., 2007), whereas Pvf1 is a ligand for the receptor tyrosine kinase (RTK) Pvr in the CZ to trigger a signaling cascade known as equilibrium signaling (Mondal et al., 2011). Together, these signaling mechanisms maintain the progenitor cell population.

RTK signaling is induced by ligand binding, but the duration and extent of the signal is regulated by membrane trafficking (Miaczynska, 2013). Membrane trafficking modulates RTK signaling by affecting receptor internalization, by regulation of the balance between receptor recycling and degradation, as well as by compartmentalization of signals (Miaczynska, 2013). Many small GTPases are known to orchestrate membrane trafficking events (Takai, Sasaki, & Matozaki, 2001). Among these, the small GTPase Ral is at the cross-roads of several membrane trafficking processes, including exocytosis, endocytosis and autophagy (Gentry, Martin, Reiner, & Der, 2014). Mammals have two Ral genes, RALA and RALB, whereas there is a single Rala gene in flies. The fly Rala protein is closely related to both human RalA (72% identity) and RalB (71% identity) (Gentry et al., 2014).

Like other small GTPases, Ral activation is regulated by specific guanine nucleotide exchange factors (GEFs) and GTPase activating proteins (GAPs). In mammals there are two main classes of RalGEFs, namely, RalGDS, Rgl1, Rgl2 and Rgl3, which contain a Ras Activation (RA)-domain for direct interaction with active Ras (Ferro & Trabalzini, 2010), and RalGPS1 and RalGPS2, which harbor a pleckstrin homology (PH) domain that interacts with phosphoinositides (Quilliam, 2006). The single RalGAP is a heterodimer consisting of an α-(RalGAPα1 or RalGAPα2 in mammals) and a β-subunit (RalGAPβ) (X. W. Chen et al., 2011; Shirakawa et al., 2009). The D. melanogaster genome comprises two genes encoding RalGEFs (FB2020_02; Thurmond et al., 2018): Rgl and the putative RalGEF CG5522 (homologous to mammalian RalGPS and thus referred to as RalGPS hereafter) (Fig. 1A). It also comprises genes homologous to those encoding the mammalian RalGAP subunits: one α-(CG5521) and one β-subunit (CG34408) (FB2020_02; Thurmond et al., 2018). The main Ral effectors that interact with GTP-bound Ral are Exo84 and Sec5 of the exocyst complex and Rlip/RalBP1 (Gentry et al., 2014). Through its interaction partners, Ral participates in exocytosis, autophagy, actin cytoskeleton dynamics, endocytosis and gene expression (Gentry et al., 2014).

**Figure 1.**
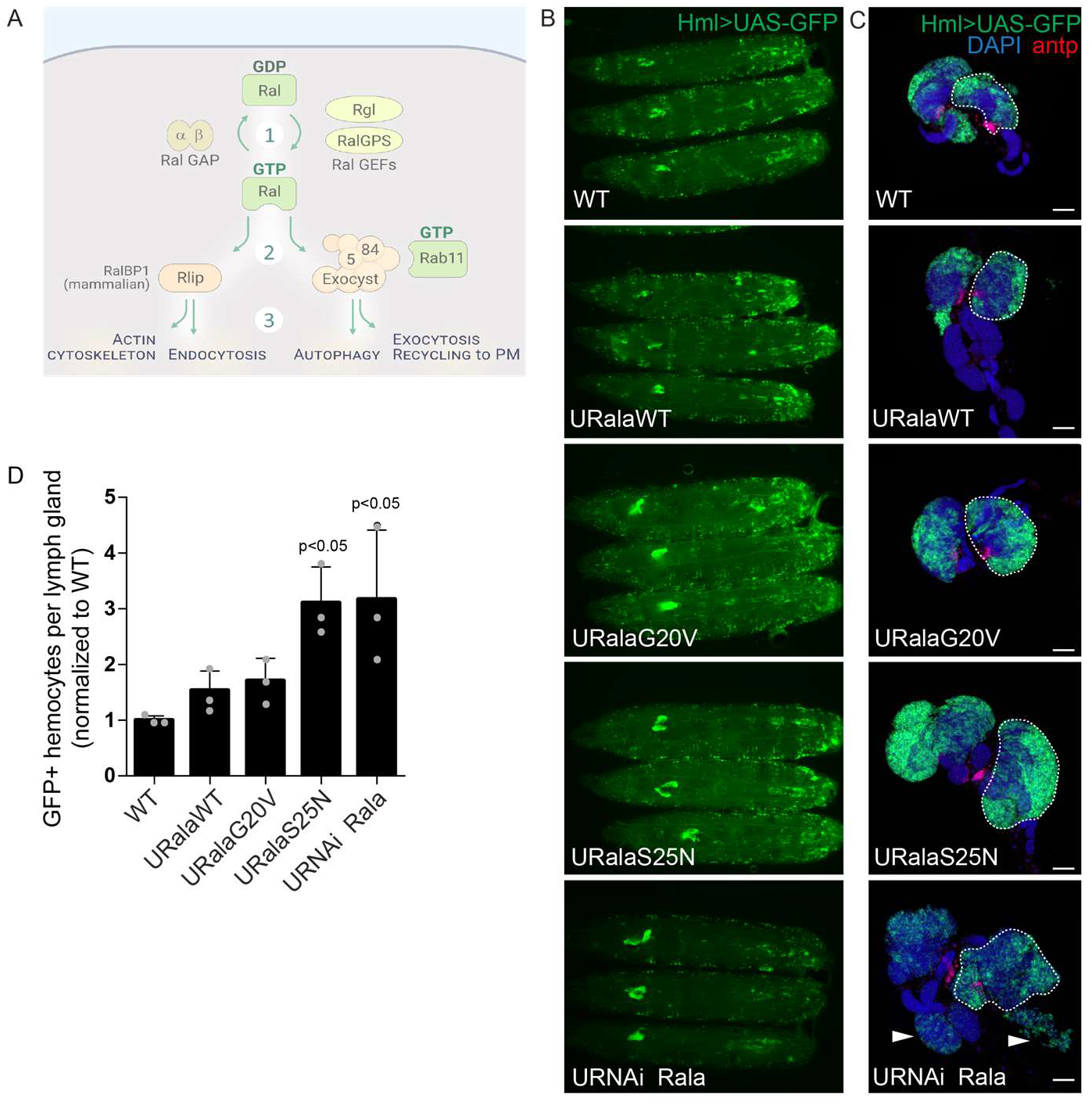
Perturbation of Rala activation in the hematopoietic system increases the number of Hml>GFP positive hemocytes in the lymph gland. **A.** Schematic overview over Rala signaling based on literature on *Drosophila* and mammalian Ral proteins. Note that RalGPS (CG5522) and the RalGAP complex (CG5521 and CG34408) have not yet been studied in flies. **B.** Wandering third instar larvae expressing wild-type (RalaWT), constitutive active (RalaG20V) or dominant negative (RalaS25N) Rala or dsRNA against Rala driven by Hml-Gal4 were imaged by widefield fluorescence microscopy to assess the hematopoietic system marked by GFP expression. **C.** Representative widefield fluorescence images of lymph glands where the cortical zone is marked by Hml-driven GFP and the PSC by antp immunostaining. One primary lobe is outlined in each image. Arrowheads point to differentiating secondary lobes. Scale bars 50 μm. **D.** Quantification of the number of GFP positive hemocytes per larval lymph gland. Graph shows average GFP-positive hemocytes per lymph gland normalized to control (WT) +/− stdev from 3 independent experiments. Statistically significant differences relative to WT are indicated (one-way ANOVA).

While there is currently no evidence that a Ras-RalGEF-Ral axis exists in flies, Rala has been found to act downstream of Rap1 (Carmena, Makarova, & Speicher, 2011; Mirey et al., 2003), another Ras-like small GTPase primarily known to control cell adhesion (Frische & Zwartkruis, 2010). Rala has been studied in embryonal development (Holly, Mavor, Zuo, & Blankenship, 2015) and the development of some fly tissues including eyes, bristles and wings (Cho & Fischer, 2011; Mirey et al., 2003; Sawamoto et al., 1999), but much is yet to be learned about the regulation, function and signaling of Rala in flies.

In this study, we investigated the role of Rala in the larval hematopoietic system as a tool to address the involvement of membrane trafficking in lymph gland homeostasis. We found that loss of *Rala* activity phenocopies *Pvr* depletion and results in hemocyte progenitor differentiation. We mapped the defect to a novel RalGPS-Rala-exocyst-Rab11 signaling axis involved in lymph gland homeostasis impinging on Pvr signaling. Impairment of this axis leads to abnormal accumulation of Pvr, suggesting a trafficking defect. Consistently, lymph gland enlargement by *Rala* loss of function is suppressed by providing constitutive STAT activity, which normally mediates the progenitor maintenance signaling from Pvr. Taken together, this places membrane trafficking as another level of regulation of hematopoietic progenitor maintenance.

## Results

### Perturbation of Rala activity in the hematopoietic system affects lymph gland homeostasis

To study the role of Rala in the hematopoietic system of *D. melanogaster* larvae, we expressed wild-type (WT), constitutively active (G20V), or dominant negative (S25N) Rala, or RNAi against Rala in the larval hematopoietic system marked by expression of GFP using the *hml*^Δ^-Gal4 driver. The resulting GFP positive cells in the lymph gland define the cortical zone of the gland, and generally correspond to differentiated plasmatocytes (Jung, Evans, Uemura, & Banerjee, 2005). Expression of constitutively active Rala in the cortical zone of the lymph gland marginally increased the number of GFP-positive cells per lymph gland (Fig. 1B-D), whereas expression of dominant negative Rala or RNAi against Rala led to a 3-fold increase in the number of GFP-positive cells per lymph gland (Fig. 1B-D), where the cortical zone was enlarged and hemocytes in the secondary lobes also started expressing GFP. Expression of RalaWT, G20V or S25N in the cortical zone of the lymph gland and in circulating hemocytes was verified by immunostaining, qPCR and western blotting (Fig. S1A-D). Immunostaining of endogenous Rala was not clear by any antibodies tested, but depletion of Rala was verified by qPCR from lymph glands (Fig. S1E) and western blotting of fat body (Fig. S1F). We also determined that *Rala* transcripts are expressed to similar levels as *rolled/mapk* mRNAs by transcriptome analysis of entire lymph glands (Table S1).

As a control, we verified that the increase in GFP-positive hemocytes in the lymph gland induced by expression of dominant negative RalaS25N could be reversed by co-expression of wildtype Rala (Fig. S2A-C). Similar to dominant negative versions of other small GTPases, dominant negative RalaS25N is thought to titrate out the RalGEF and thereby inhibit the function of endogenous Rala (Mirey et al., 2003; Powers, O’Neill, & Wigler, 1989).Co-expression of wild-type Rala probably allows enough Rala molecules to become loaded with GTP due to the high cellular GTP concentration and thereby perform their endogenous function. Finally, the increase in GFP-positive hemocytes in the lymph gland upon depletion of Rala with an shRNA targeting the 3’UTR could be partially rescued by expression of wild-type Rala (Fig. S2D-F).

In terms of development into differentiated hemocytes, expression of RalaWT, RalaG20V or RalaS25N in the cortical zone did not affect the percentage of Hnt-positive crystal cells per primary lobe (5-8% per lobe), whereas fewer crystal cells were observed upon Rala depletion (2.6% per primary lobe) (Fig. S3A). Similarly, no lamellocytes were detected in circulation upon expression of RalaWT, RalaG20V or RalaS25N, but were sometimes observed upon Rala depletion (Fig. S3B-B’). Immunostaining against P1 (Nimrod C1 antigen) showed that the Hml-positive hemocytes in all genotypes were plasmatocytes, both in circulation and in the lymph gland (Fig. S3C, D). Finally, Rala-depleted hemocytes were competent at phagocytosis (Fig. S3E). Taken together, this indicates that perturbation of Rala’s function in the cortical zone of the lymph gland disturbs lymph gland homeostasis and leads to increased numbers of differentiated cells in the cortical zone.

### Ras and Rala act independently on hemocytes

Since active Ras has previously been shown to strongly induce proliferation of circulating hemocytes in *Drosophila* (Asha et al., 2003; Zettervall et al., 2004) and Rala is activated downstream of active Ras in mammals, we investigated the effect of Ras and Rala on circulating hemocytes in 3^rd^ instar larvae. In line with previous reports (Asha et al., 2003), expression of active RasV12 in the larval hematopoietic system with the *hml*^Δ^-Gal4 driver increased hemocyte numbers by 50-100 fold (Fig. 2A, C) and induced differentiation of lamellocytes, observed as actin-rich large cells (Fig. S4A). As Ras transmits signals through multiple effector pathways (Cox & Der, 2010), we expressed effector loop mutations of Ras in an attempt to define the Ras effector(s) involved. These effector loop mutations, namely, S35, G37, and C40, have been shown in mammalian cells to maintain interaction with only one of the three main Ras effectors, that is, Raf, RalGDS or PI3K, respectively (Joneson, White, Wigler, & Bar-Sagi, 1996; White, Vale, Camonis, Schaefer, & Wigler, 1996) (Fig. 2B). When expressed in the larval hematopoietic system, RasV12_S35 enhanced hemocyte numbers similar to RasV12 itself, whereas RasV12_G37 and RasV12_C40 were significantly less effective (Fig. 2A, C and S4A). These results were not merely related to differences in expression levels (Fig. S4B). This suggested that the RasV12-induced hemocyte proliferation primarily depends on Raf-MAPK signaling. Indeed, when MAPK was depleted in hemocytes expressing RasV12, hemocyte numbers returned to close to wild-type levels (Fig. 2D and S4C). However, when Rala was depleted in hemocytes expressing RasV12, hemocyte numbers surprisingly increased even further, compared to RasV12 alone (Fig. 2D and S4D). This suggests not only that Rala does not mediate Ras signals, but that it actually opposes Ras in this process. Moreover, depletion of MAPK in hemocytes expressing RasV12_S35 (activating the Raf-MEK-MAPK signaling axis) abolished the RasV12_S35-induced hemocyte proliferation (Fig. S4C). Rala depletion in hemocytes expressing RasV12_G37 (supposedly activating RalGEF-Ral signaling) did not affect the number of hemocytes (Fig. S4E). In fact, depleting MAPK in these RasV12_G37-expressing hemocytes strongly inhibited the increased hemocyte numbers (Fig. S4C). It thus appears that RasV12_G37 allele in this experimental paradigm may not signal through Rala, but mainly through the MAPK pathway. In conclusion, it appears that the RasV12_G37 effector loop mutant cannot be used to study Rala signaling in the hematopoietic system of flies.

**Figure 2.**
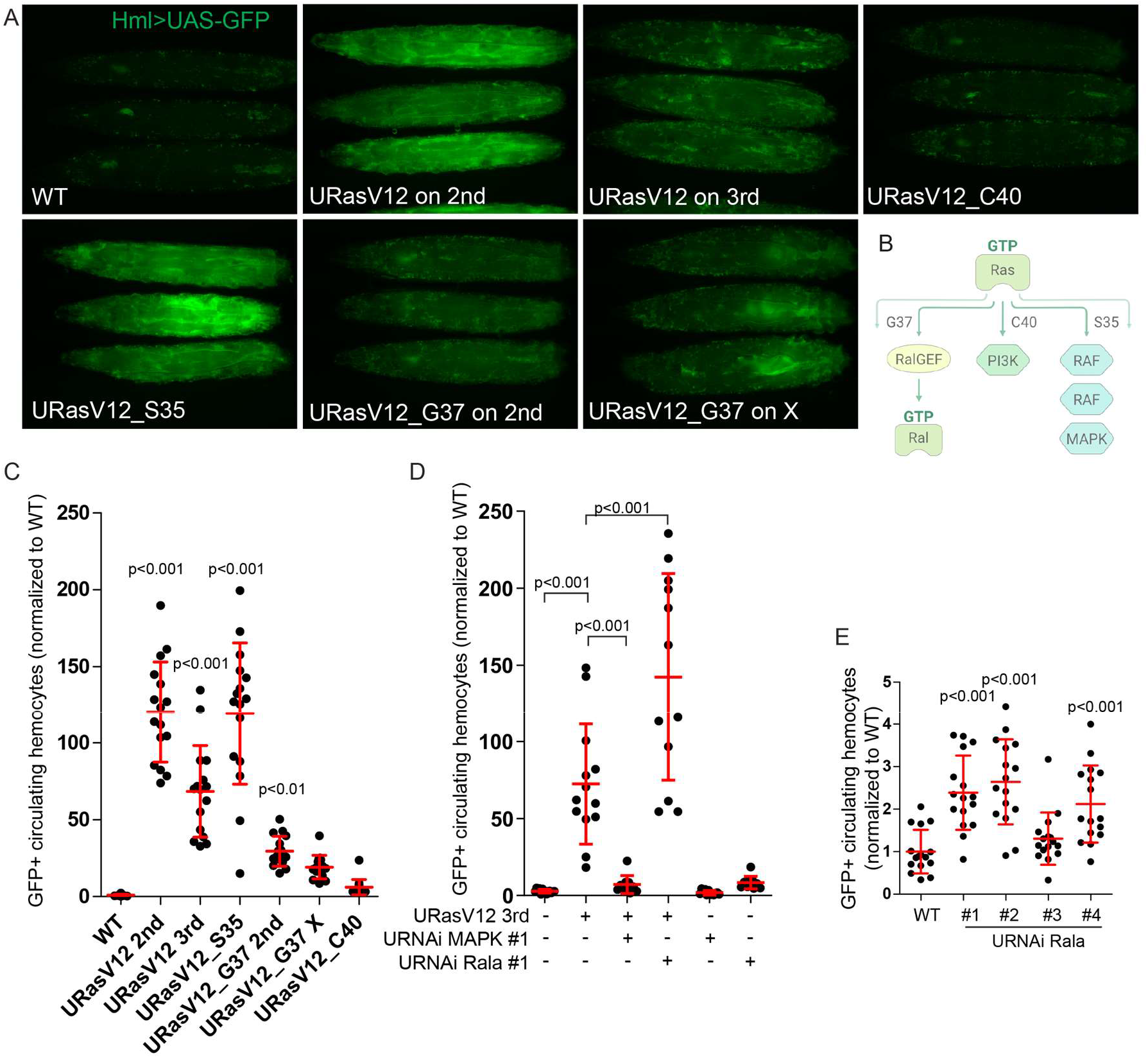
RasV12-induced hemocyte overproliferation does not depend on Rala. **A.** Wandering third instar larvae expressing RasV12, RasV12_S35, RasV12_C40 or RasV12_G37 driven by Hml-Gal4 were imaged by widefield fluorescence microscopy to assess the hematopoietic system marked by GFP expression. **B.** Schematic overview over the three main effectors downstream of Ras. **C.** Larvae as shown in A were opened to release the hemolymph. The number of GFP-positive circulating hemocytes was quantified by high-content microscopy. The graph shows mean +/− stdev from a total of 12 larvae from 4 independent experiments. **D.-E.** GFP-positive circulating hemocytes were quantified as in C. The graph shows mean (normalized to control, WT) +/− stdev from a total of 12 larvae from 4 independent experiments (**D**) or a total of 16 larvae in 3 independent experiments (**E**). Statistically significant differences in C-E are indicated (one-way ANOVA).

To obtain further evidence that Rala regulates the number of circulating hemocytes, we depleted Rala using RNAi. This resulted in a 2.5-fold increase in hemocyte numbers by 3 out of 4 RNAi lines tested (Fig. 2E). Taken together, these data indicate that RasV12-induced hemocyte proliferation is mediated by MAPK, that Rala signaling occurs independently of Ras, and that Rala opposes Ras-MAPK-dependent hemocyte proliferation.

### Rala is activated by the GEF RalGPS in hemocytes

To gain a better understanding of signaling upstream of Rala in *Drosophila* hemocytes, we investigated the activation of Rala at the biochemical level. To this end, we used S2 cells as a cell culture model of hemocytes of embryonic origin (Cherbas et al., 2011). To detect GTP-bound active Rala, we used recombinant GST-tagged Ral-binding domains (RBDs) of either the human or the *Drosophila* versions of the Ral effectors Rlip/RalBP1 and Sec5 (Fig. S5A). Each GST-RBD protein strongly interacted with 3xFlag-tagged constitutively active GTP-locked RalaG20V expressed in S2 cells, and also interacted with 3xFlag-RalaWT, albeit to a much weaker degree (Fig. S5B). Expression of WT or constitutively active Rala did not affect PI3K-Akt or Raf-MAPK signaling as no significant changes in phosphorylation of Akt or MAPK were detected (Fig. S5B).

The *D. melanogaster* genome encodes two RalGEFs: Rgl and CG5522 (homologous to mammalian RalGPS and thus referred to as RalGPS hereafter). It also encodes one α-(CG5521) and one β-subunit (CG34408) of the RalGAP heterodimer. To assess the impact of these proteins on Rala activation, we first demonstrated that GST-Rlip-RBD could indeed pull down endogenous GTP-loaded Rala (Fig. 3A, lanes 3-5) and confirmed that dsRNAs targeting the *Drosophila RalGEF* and *RalGAP* transcripts reduced their respective expression levels as measured by qPCR (Fig. S5C). Markedly, depletion of RalGPS strongly decreased the levels of active Rala (Fig. 3A, lanes 6-7), whereas Rgl depletion had no effect (Fig. 3A, lanes 8-10). Conversely, depletion of the RalGAP α- or β-subunit strongly increased active Rala levels (Fig. 3A, lanes 11-13).

**Figure 3.**
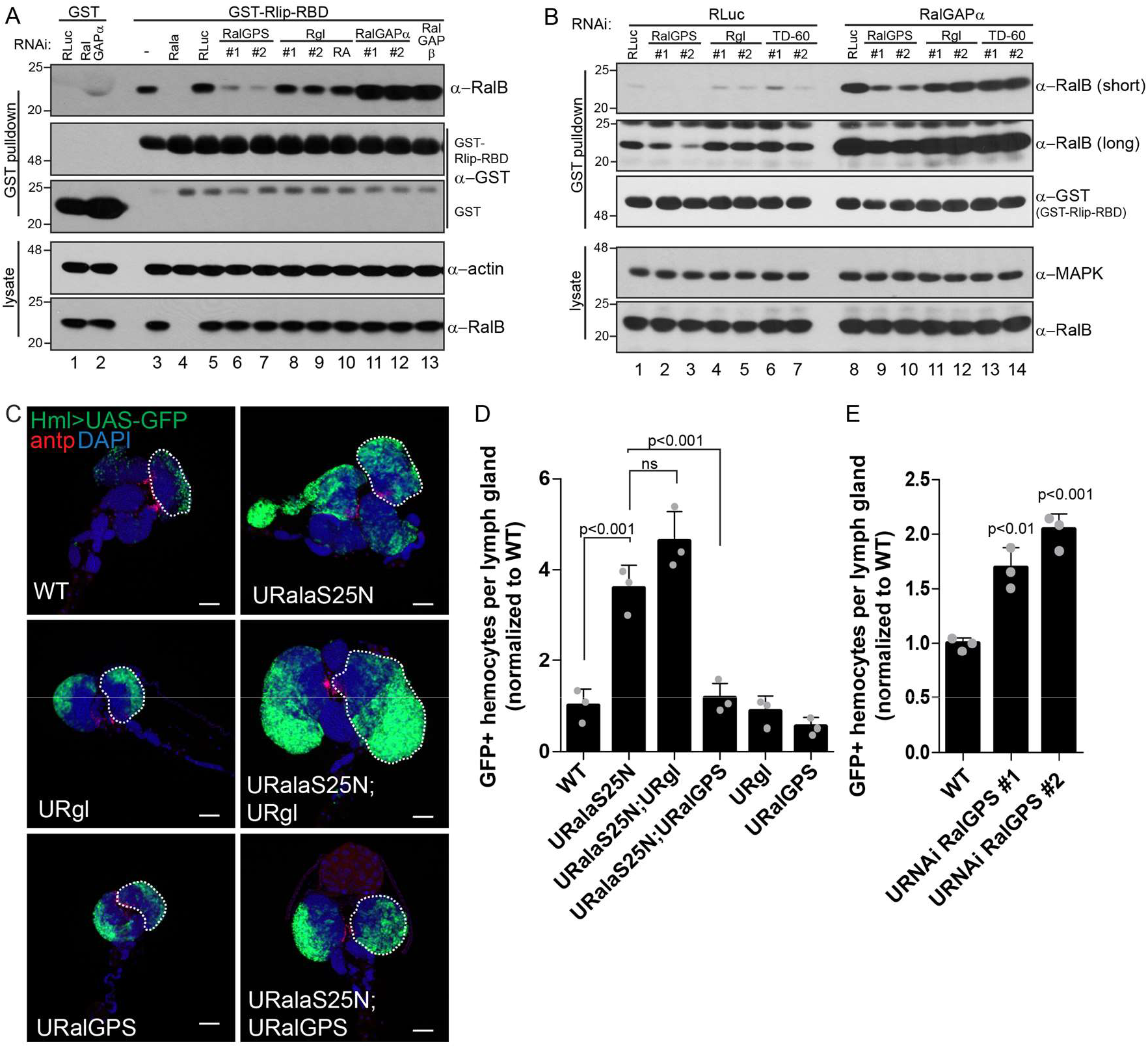
Expression of RalGPS and not Rgl1 rescues the lymph gland hyperplasia induced by dominant negative Rala. **A-B.** S2 cells were treated with dsRNA against the indicated targets. Recombinant GST or GST-Rlip-RBD bound to GSH sepharose was incubated with the S2 cell lysates and the resulting pull-downs analyzed by immunoblotting performed with the indicated antibodies. **C.** Representative widefield fluorescence images of lymph glands expressing Hml-driven dominant negative Rala (RalaS25N) alone or in combination with the RalGEFs Rgl or RalGPS. The cortical zone is marked by Hml-driven GFP and the PSC by antp immunofluorescence. One primary lobe is outlined in each image. Scale bar 50 μm. **D-E.** Quantification of the number of GFP positive hemocytes per larval lymph gland. Graphs show average GFP-positive hemocytes per lymph gland normalized to control (WT) +/− stdev from 3 independent experiments. Statistically significant differences are indicated (one-way ANOVA). ns: non-significant.

We next evaluated the impact of co-depleting RalGAPα along with the distinct RalGEFs. Recently, TD-60 (also known as RCC1) was described as a novel GEF for mammalian Ral (Papini et al., 2015). We tested whether the *Drosophila* homolog CG9135 (referred to as TD-60 hereafter) was required for Rala activity. We knocked down TD-60 either alone or in combination with of RalGAPα, but found that only depletion of RalGPS resulted in abrogation of Rala activity (Fig. 3B and Fig. S5D). We conclude that RalGPS is the GEF that activates Rala in S2 cells at steady state and that Rala activity is tightly controlled by the RalGAP heterodimer.

Since we observed no evidence of Rala mediating the hemocyte overproliferation observed upon RasV12 expression (Fig. 2 and S4), we next asked whether we could biochemically detect Rala activation downstream of active Ras in S2 cells. To this end, we used three different strategies: 1) stimulation of *Drosophila* EGFR (DER)-expressing S2 cells with supernatant from cells producing the EGFR ligand Spitz (Fig. S6A), 2) induction by heat-shock of a constitutively active Sevenless receptor (Sev-^S11^) (Basler, Christen, & Hafen, 1991; Therrien, Wong, & Rubin, 1998) (Fig. S6B) and 3) stimulation of S2 cells with human recombinant insulin over a time-course of 2.5 to 80 minutes (Fig. S6C). In each case, activation of Ras-MAPK signaling was confirmed by phosphorylation of MAPK, whereas activation of Rala was assessed by GST-RBD pulldown as described above. In neither of these three experimental setups did we detect any activation of Rala in the time-frame tested. We conclude that *Drosophila* Rala is not activated downstream of active Ras upon any of these stimuli.

Since we found RalGPS and not Rgl to be the GEF for Rala in unstimulated S2 cells, we wondered which RalGEF was mediating activation of Rala in the lymph gland. We detected expression of both GEFs by transcriptome analysis of entire lymph glands, but RalGPS mRNA was about 4 times more abundant than Rgl mRNA (Table S1). Interestingly, in contrast to Rgl, we found that RalGPS overexpression completely suppressed the increased number of GFP-positive hemocytes caused by RalaS25N expression in the cortical zone (Fig. 3C-D). Overexpression of either Rgl or RalGPS alone did not significantly affect the number of GFP-positive hemocytes per lymph gland (Fig. 3C-D). Conversely, reduction of RalGPS by RNAi phenocopied a loss of Rala activity and led to an increased number of GFP-positive hemocytes per lymph gland (Fig. 3E, S7A). We conclude that RalGPS is the GEF responsible for Rala activation in the lymph gland, similar to the situation in S2 cells.

### The exocyst complex and Rab11 act downstream of Rala in hemocytes

To further dissect the Rala signaling axis active in the lymph gland, we separately depleted by RNAi the three main effectors of Rala, namely, Sec5 and Exo84 of the exocyst complex and Rlip/RalBP1. Depletion of Sec5 or Exo84 led to an increase in GFP-positive hemocyte numbers similar to Rala depletion, whereas depletion of Rlip had no effect (Fig. 4A-B and S7B and C). Since exocyst components can exist in different sub-complexes and have functions independent of the full exocyst complex (Wu & Guo, 2015), we individually depleted most of the 8 exocyst subunits and found that depletion of either Sec6, Sec8 or Sec15 led to enlarged cortical zones, similar to the depletion of Exo84 or Sec5 (Fig. S7D). It thus appears that the exocyst complex, and not Rlip/RalBP1, functions downstream of Rala for lymph gland homeostasis.

**Figure 4.**
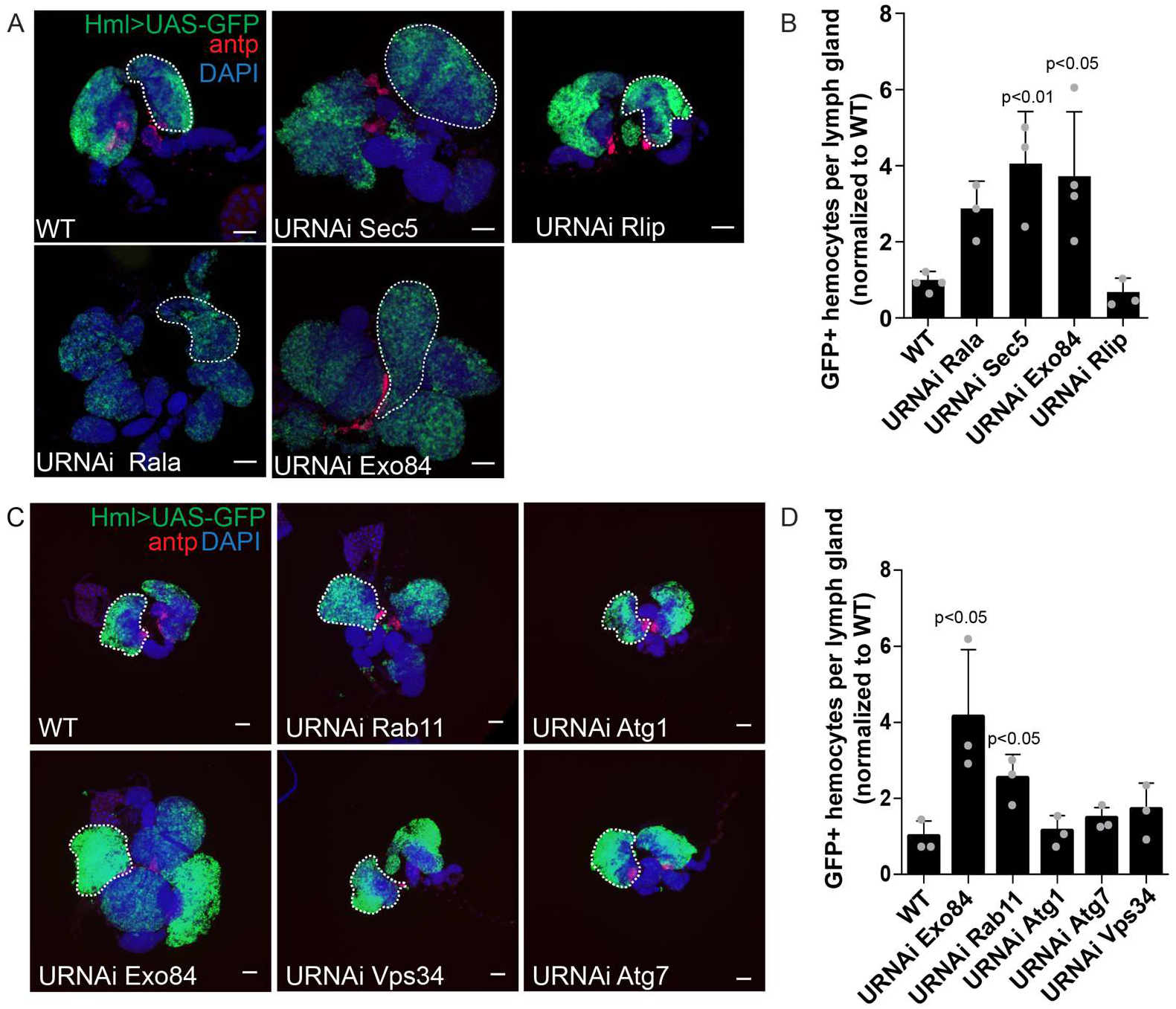
Depletion of exocyst components or Rab11 and not Rlip or components of the autophagy machinery induces enlargement of the lymph gland. **A and C.** Representative widefield fluorescence images of lymph glands expressing dsRNA against Rala, Sec5, Exo84 or Rlip (A), Rab11, Vps34, Atg1 or Atg7 (B) driven by Hml-Gal4. The cortical zone is marked by Hml-driven GFP and the PSC by antp immunofluorescence. One primary lobe is outlined in each image. Scale bar 50 μm. **B and D.** Quantification of the number of GFP positive hemocytes per larval lymph gland. Graphs show average GFP-positive hemocytes per lymph gland normalized to control (WT) +/− stdev from 3 independent experiments. Statistically significant differences are indicated (one-way ANOVA).

The exocyst complex is involved in various membrane trafficking events such as exocytosis, endosome / membrane receptor recycling and autophagy (Wu & Guo, 2015). To pinpoint which of these functions might be involved in lymph gland homeostasis, we depleted either the small GTPase Rab11 (a central player in vesicular endocytosis and recycling, and a direct interaction partner of the exocyst complex) or several components of the autophagy machinery. Depletion of Rab11 led to an increase in the number of GFP-positive hemocytes per lymph gland, whereas depletion of various autophagy components had no effect (Fig. 4C-D and S8A-B). These findings suggest that membrane trafficking events linked to endocytosis and/or endosome recycling are important for lymph gland homeostasis.

The RTK Pvr controls the differentiation of hemocytes from prohemocytes. Increased Pvr activity in the cortical zone hinders differentiation of prohemocytes from the medullary zone, resulting in small lymph glands. Conversely, Pvr depletion triggers hemocyte differentiation from the medullary zone leading to enlarged cortical zones (Fig. S9A-B) (Mondal et al., 2011). We stained for Pvr, expecting to find decreased levels in the cortical zone of Rala or exocyst depleted lymph glands. To our surprise, Pvr levels were increased upon depletion of Rala, Sec5, Exo84 or Rab11 (Fig. 5A). Imaging of a smaller area of the cortical zone showed that Pvr appears to accumulate close to the plasma membrane when Rala or Sec5 are depleted (Fig. 5B).

**Figure 5.**
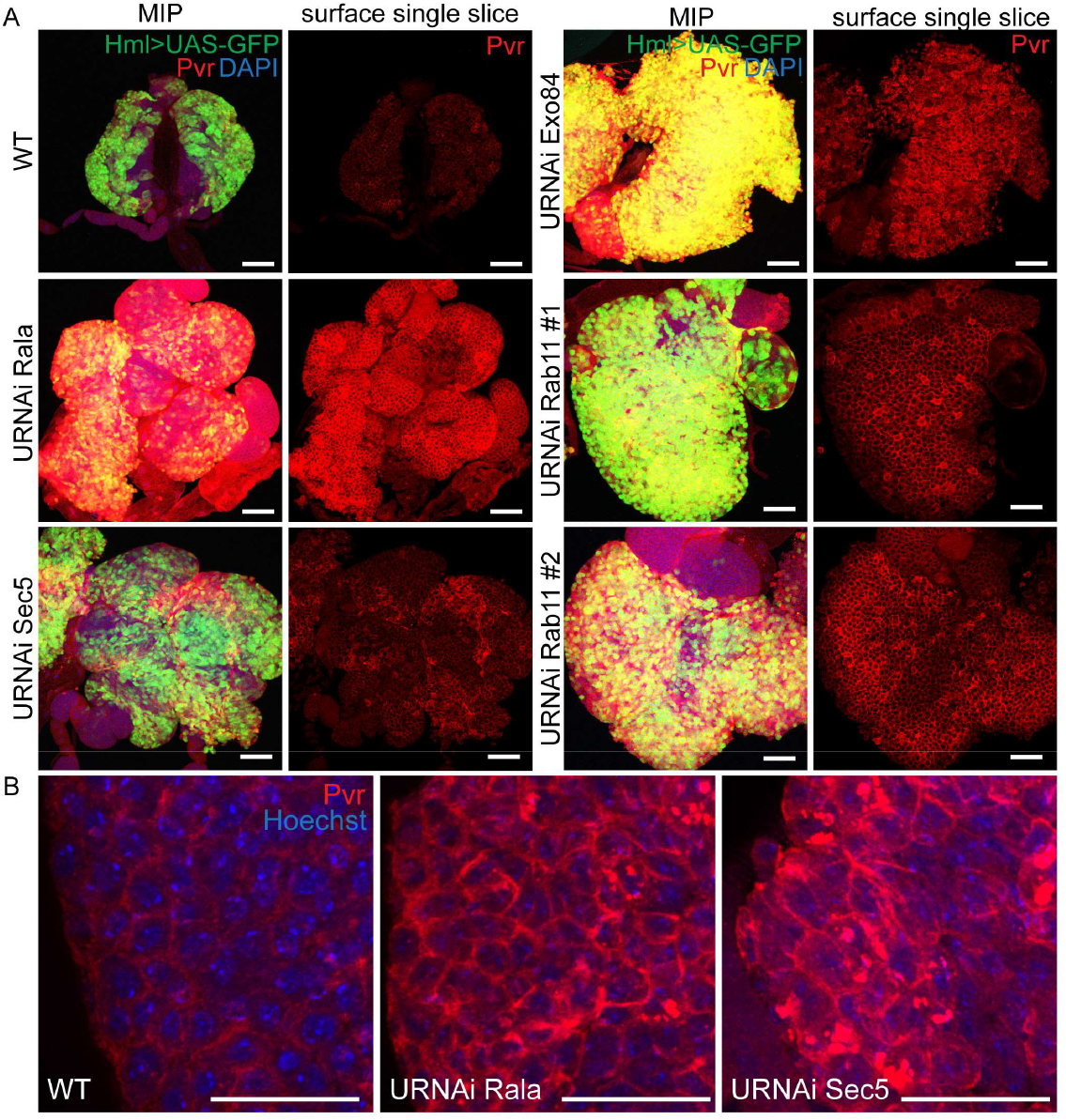
Pvr accumulates upon Rala or exocyst depletion. **A.** Maximum intensity projections (MIP) or surface single slices of confocal micrographs of immunostaining against Pvr in lymph glands expressing dsRNA against Rala, Sec5, Exo84 or Rab11 driven by Hml-Gal4. Scale bar 50 μm. **B.** Confocal micrographs of an area of the cortical zone of lymph glands expressing dsRNA against Rala or Sec5 driven by Hml-Gal4 compared to WT. MIPs of 10 slices closest to the surface from confocal imaging of immunostaining against Pvr. Scale bars 20 μm. All images for each subpanel were captured with identical settings below pixel value saturation and post-processed identically.

We currently do not know the mechanism affected by the loss of Rala / exocyst activity, which impinges on Pvr levels at the plasma membrane. Nevertheless, since the loss of Rala / exocyst activity in the cortical zone phenocopied Pvr impairment, we conclude that the elevated levels of Pvr proteins observed upon Rala / exocyst depletion have also reduced activity. To verify this possibility, we ectopically expressed Pvr in Rala or exocyst depleted cortical zones. As shown in Fig. 6A-B, Pvr overexpression in the cortical zone in combination with depletion of either Rala, Sec5, Exo84 or Rab11 rescued the enlarged lymph gland size (Fig. 6A-B). Pvr expression could also rescue the enlarged cortical zone induced by dominant negative RalaS25N (Fig. S9C-D). We assume that upon overexpression of Pvr, enough productive signaling occurs and thereby restore lymph gland homeostasis. It is known that Pvr equilibrium signaling is mediated mainly through Stat92E (Mondal et al., 2011). Consistently, expression of a dominant active Stat92E, Stat92EΔNΔC (Ekas et al., 2010), rescued the enlargement of the cortical zone induced by Rala depletion (Fig. 6C-D).

**Figure 6.**
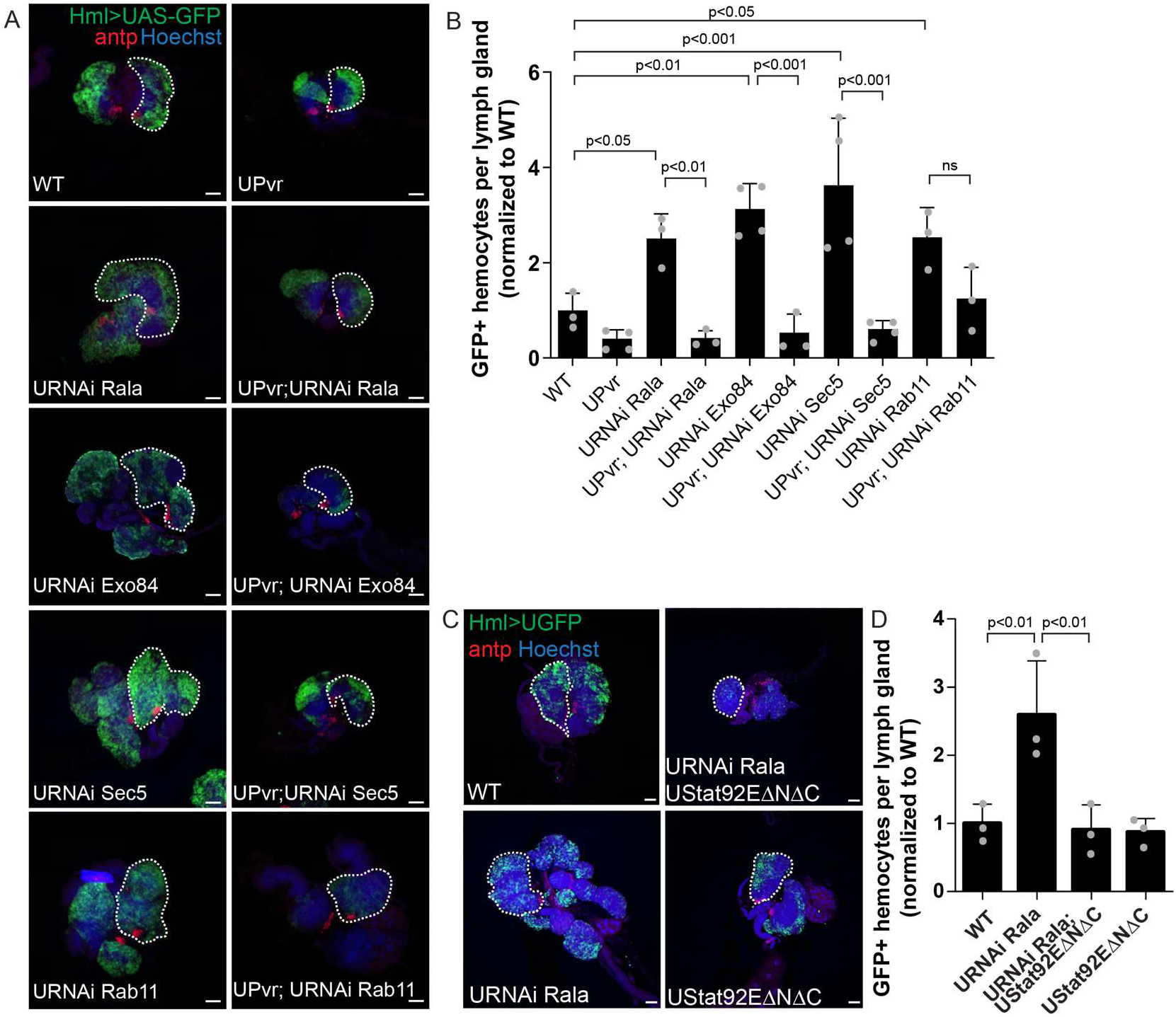
Expression of Pvr or Stat92E-GOF rescues lymph gland homeostasis. **A and C.** Representative widefield fluorescence images of lymph glands expressing dsRNA against Rala, Exo84, Sec5 or Rab11 alone or in combination with Pvr expression (A) or in combination with Stat92EΔNΔC (Stat92E-GOF) (C) driven by Hml-Gal4. The cortical zone is marked by Hml-driven GFP and the PSC by antp immunofluorescence. One primary lobe is outlined in each image. Scale bars 50 μm. **B and D.** Quantification of the number of GFP positive hemocytes per larval lymph gland. Graph shows average GFP-positive hemocytes per lymph gland normalized to control (WT) +/− stdev from 3 independent experiments. Statistically significant differences are indicated (one-way ANOVA).

In summary, we propose a model where a RalGPS-Rala-exocyst-Rab11 axis is required for lymph gland homeostasis (Fig. 7). In the absence of a functional Rala axis, there is a loss of equilibrium signaling and a loss of feedback within the lymph gland, resulting in the expansion of the cortical zone. We suggest that the loss of equilibrium signaling is due to the RalGPS-Rala-exocyst-Rab11 axis normally regulating a step impinging on Pvr accumulation at the plasma membrane and required for downstream signaling.

**Figure 7.**
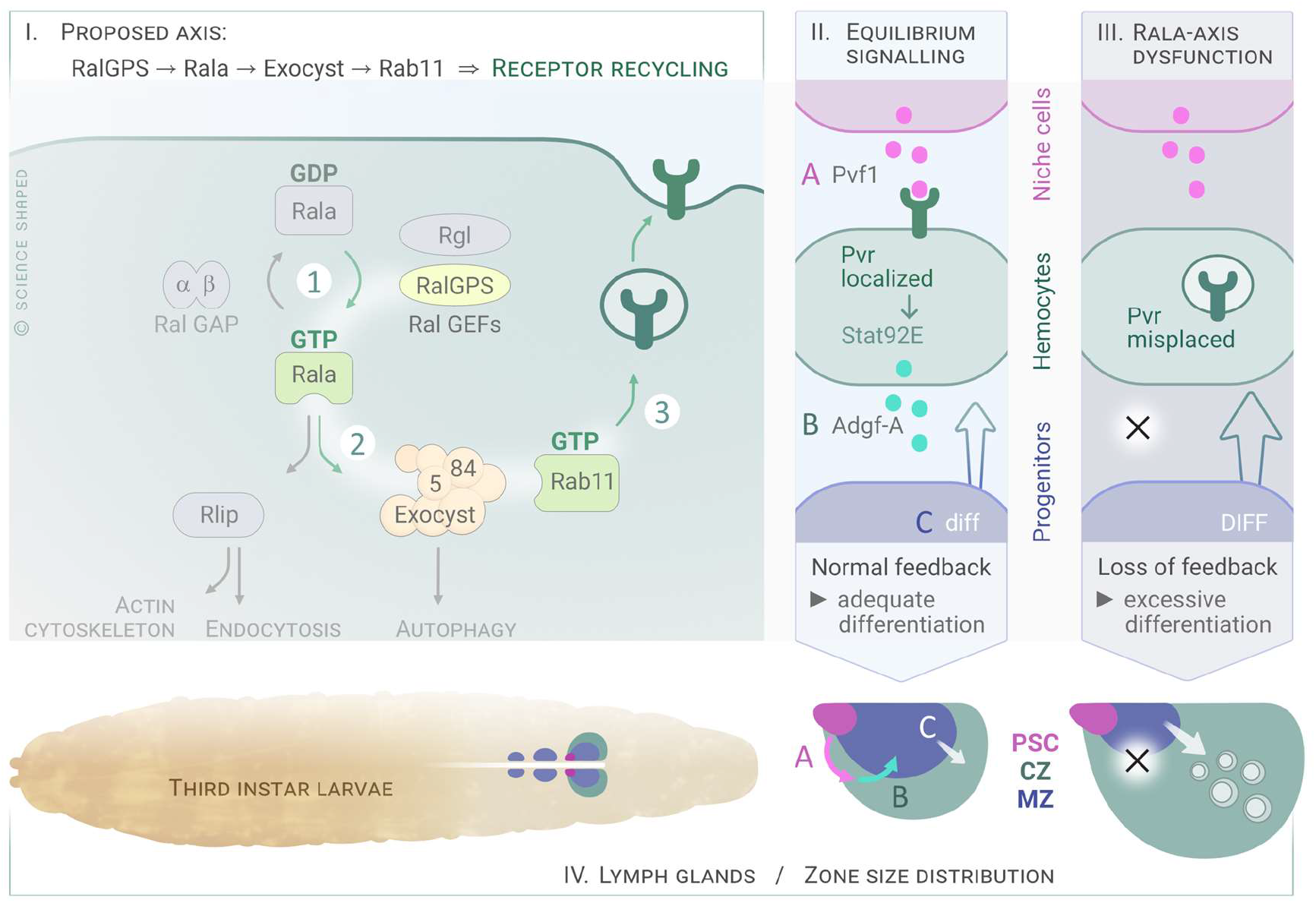
The RalGPS-Rala-Exocyst-Rab11 axis regulates lymph gland homeostasis through effects on Pvr. To summarize our findings, we propose the following model: I. The proposed Rala axis necessary for lymph gland homeostasis consists of RalGPS, Rala, the exocyst complex and Rab11. RalGPS is the GEF responsible for activating Rala in the cortical zone (1). Downstream of Rala, the exocyst complex (2) and Rab11 (3) are responsible for correct localization of Pvr. II. In wild-type larvae the equilibrium signal is intact leading to normal feedback to the cortical zone and adequate differentiation of hemocytes. III. When the Rala axis is dysfunctional, the equilibrium signal and feedback is lost, leading to an excessive expansion of the cortical zone (IV).

## Discussion

In contrast to mammalian VEGF and PDGF receptors, which encompass multiple isoforms, *Drosophila* encodes a single PDGF/VEGF-like receptor, known as Pvr. In the developing lymph gland, Pvf-induced Pvr signaling occurs in differentiating hemocytes of the cortical zone. This event regulates progenitor maintenance in the adjacent medullary zone through a process called “equilibrium signaling”, which involves STAT92E activity as well as ADGF-A expression downstream of Pvr (Mondal et al., 2011). ADGF-A is a secreted enzyme that keeps in check the levels of extracellular adenosine (Zurovec, Dolezal, Gazi, Pavlova, & Bryant, 2002), an inducer of progenitor differentiation (Mondal et al., 2011). A number of factors such as Bip1, Nup98, RpS8, and Sd have been shown to impinge on Pvr expression and thereby modulate equilibrium signaling (Ferguson & Martinez-Agosto, 2017; Mondal, Shim, Evans, & Banerjee, 2014). Although their respective mechanisms have yet to be determined, none appeared to affect Pvr trafficking. In the present work, we described a novel Rala-exocyst-Rab11 signaling axis as an additional level of regulation of Pvr signaling through receptor trafficking that accordingly impinges on equilibrium signaling.

We found that depletion of Rala, exocyst or Rab11 in the cortical zone of the lymph gland leads to hyperplasia of this tissue. Presumably, this is because Pvr is incorrectly trafficked, which impedes its normal signaling, disrupts the equilibrium signal and results in excessive progenitor differentiation. Endocytosis and recycling strongly affect signaling from receptors of the mammalian PDGFR and VEGFR families (Hellberg, Schmees, Karlsson, Ahgren, & Heldin, 2009; Horowitz & Seerapu, 2012; Kawada et al., 2009; Lennartsson, Wardega, Engström, Hellman, & Heldin, 2006; Nakayama et al., 2013). PDGF and VEGF receptor signaling is also regulated at internal compartments, such early endosomes (Ballmer-Hofer, Andersson, Ratcliffe, & Berger, 2011; Lanahan et al., 2010; Muratoglu, Mikhailenko, Newton, Migliorini, & Strickland, 2010; Wang, Pennock, Chen, Kazlauskas, & Wang, 2004). Furthermore, Pvr localization and trafficking is important for its signaling in physiological settings in *Drosophila*, such as in border cells, which also depends on Rab11 (Janssens, Sung, & Rorth, 2010; Jekely, Sung, Luque, & Rorth, 2005). We found that loss of the Rala signaling axis leads to Pvr accumulation close to the plasma membrane. The exact localization and nature of compartment is yet to be determined, but genetic experiments point to a defect in receptor recycling. The observed increase in Pvr levels might reflect a reduction in receptor downregulation due to trafficking defects, or an upregulation of Pvr expression as a compensatory mechanism to cope with the perturbed trafficking and impeded signaling, or a combination of both. Still, exogenous expression of Pvr can rescue loss of Rala function. We attribute this to correct localization for signaling of newly synthesized receptors, following a different trafficking path than existing receptors destined for recycling or downregulation, possibly also in combination with exogenous expression simply allowing enough receptor molecules to signal correctly. Rala and the exocyst-Rab11 signaling axis therefore represent another level to allow regulation of Pvr signaling through membrane trafficking.

Previous studies have linked perturbations of endocytic trafficking to changes in levels of differentiated hemocytes (Rohan J. Khadilkar et al., 2017; R. J. Khadilkar et al., 2014; Kim et al., 2017; Korolchuk et al., 2007; Kulkarni, Khadilkar, Magadi, & Inamdar, 2011). Interestingly, loss of the endocytic protein Asrij or the small GTPase Arf79f, which has an essential function in vesicular trafficking, results in loss of prohemocyte maintenance and premature differentiation (R. J. Khadilkar et al., 2014; Kulkarni et al., 2011). However, in this case, Asrij and Arf79f were found to function downstream of Pvr (R. J. Khadilkar et al., 2014), indicating that membrane trafficking is important at multiple levels of signaling for progenitor maintenance. Future studies should address the connection between the ARF1-Asrij and the Rala-exocyst axis in maintenance of blood cell progenitors.

In mammalian cells, RalA/B are effectors downstream of Ras. As in previous tissues studied in *Drosophila* (Mirey et al., 2003), we did not find Rala to act downstream of Ras in the larval hematopoietic system or in S2 cells. Instead, we found that Rala is regulated by its GEF RalGPS in S2 cells at steady-state and in the lymph gland. In contrast to Rgl, RalGPS does not contain a RAS exchanger motif (REM) or RAS association (RA) domain, which would allow direct regulation by small GTPases like Rap1 and Ras. Instead, RalGPS contains a C-terminal PH domain that is sufficient for membrane targeting and necessary for Ral activation in mammalian cells (de Bruyn et al., 2000; Rebhun, Chen, & Quilliam, 2000). Mammalian RalGPS1/2 also contain a proline-rich sequence with PxxP motifs recognized by SH3 domain-containing proteins. Little is known about the function and regulation of RalGPS proteins. Mechanistically, human RalGPS1 and RalGPS2 localize differently and affect cytokinesis at different stages (Cascone et al., 2008). Murine RalGPS2 is involved in formation of tunneling nanotubes (D’Aloia et al., 2018) and its PH-PxxP domain promotes neurite outgrowth in cell culture by acting as a dominant negative for RalA (Ceriani, Amigoni, Scandiuzzi, Berruti, & Martegani, 2010). Regarding regulation, murine RalGPS2 appears to bind PI(4,5)P2, PI(3,4)P2, PI(3,5)P2 and PI(3,4,5)P3 (Ceriani et al., 2007). Furthermore, RalGPS2 localization was partly modified and its activation of RalA diminished by the PI3K inhibitor wortmannin (Ceriani et al., 2007). Regulation of RalGPS by PtdIns kinases and phosphatases or by SH3-domain containing proteins in the lymph gland or other fly tissues is yet to be addressed.

Depletion of either the regulatory or the catalytic subunit of the RalGAP complex strongly increased Rala activation in S2 cells (Fig. 3). However, activating Rala in the lymph gland by overexpression of RalGPS did not affect lymph gland morphology (Fig. 3C-D), although expression of constitutively active Rala slightly increased the number of Hml-GFP positive cells (Fig 1B-D). In mammalian cells, the RalGAP complex is directly regulated by the serine/threonine kinase AKT, which phosphorylates the catalytic subunit RalGAPα and inhibits its interaction with RalA (Q. Chen et al., 2014; X. W. Chen et al., 2011). Insulin stimulates this phosphorylation, to increase RalA activity and promote exocytosis of GLUT4-containing vesicles resulting in increased glucose uptake (X. W. Chen, Leto, Chiang, Wang, & Saltiel, 2007; X. W. Chen et al., 2011). We have not directly addressed the regulation of the RalGAP complex in flies, but stimulation of S2 cells with human insulin did not appear to activate Rala under the conditions tested (Fig. S3).

Although it was previously reported that Atg6 mutant flies have enlarged lymph glands (Shravage, Hill, Powers, Wu, & Baehrecke, 2013), we did not find any effect on lymph gland size of depleting autophagy components specifically in the cortical zone of the lymph gland. Nutritional signals are well established as regulating the hematopoietic system as a systemic signal, entailing cues from the brain and the fat body to hemocytes (Dragojlovic-Munther & Martinez-Agosto, 2012; Shim, Mukherjee, & Banerjee, 2012). Starved larvae show premature and excessive differentiation of hemocytes (Benmimoun, Polesello, Waltzer, & Haenlin, 2012; Shim et al., 2012) and the enlarged lymph glands in autophagy-deficient animals might be related to changes in metabolic signaling.

RalA/B is involved in several human cancer types, either dependent or independent of Ras mutations (Gentry et al., 2014). Overexpression of Ral proteins or RalGEFs or increased Ral activation has been observed in cancer cell lines and patient samples (Gentry et al., 2014). Depletion or deletion of RalA/B in cell lines and in mice reduced cancer-relevant processes, such as anchorage-independent growth, metastasis and invasion of several cancer types. These cancer-promoting effects of RalA/B have been linked to both Rlip/RalBP1 and the exocyst in a manner that depends on the Ral isoform and the cancer type (Yan & Theodorescu, 2018). Components of the exocyst have been found to be upregulated in certain cancer types and to affect cancer-relevant cell biology dependent or independent of Ral proteins (Tanaka, Goto, & Iino, 2017). The mammalian PDGFR and VEGFR families are involved in normal hematopoiesis and various blood dysplasias (Demoulin & Montano-Almendras, 2012; Gerber & Ferrara, 2003). Activating mutations in the PDGFR family member Flt3 is observed in approximately 30% of acute myeloid leukemia cases (Cancer Genome Atlas Research et al., 2013). Interestingly, this mutated receptor is partly retained in the endoplasmic reticulum, from where it drives oncogenic signaling (Choudhary et al., 2009). Future studies should address the role of RalA/B and the exocyst in normal and oncogenic signaling from RTKs in the PDGFR and VEGFR families.

## Materials and methods

### Fly stocks

*hml*^Δ^-*Gal4, UAS-2XeGFP* BL30142 and BL30140, P{w[+mC]=Cg-GAL4.A BL7011, TRiP lines UAS-RNAi Rala BL29580 and BL34375, UAS-RNAi Sec5 BL27526, UAS-RNAi Exo84 BL28712, UAS-RNAi MAPK BL36058 and BL34855, UAS-RNAi Rab11 BL27730, UAS-RNAi Atg1 BL26731, UAS RNAi Atg4 BL35740, UAS RNAi Atg5 BL27551, UAS-RNAi Atg6 BL35741, UAS-RNAi Atg7 BL27707, UAS-RNAi Atg12 BL27552, UAS-RNAi Atg16 BL28060, UAS RNAi Atg18 BL34714 were obtained from the Bloomington Stock Center. RNAi lines UAS-RNAi Rala 105296 and 43622, UAS-RNAi Sec5 28873 and 28874, UAS-RNAi RalGPS/CG5522 40595 and 40596, UAS-RNAi Exo84 108650 and 30112, UAS-RNAi Rlip 16244 and 101635, UAS-RNAi MAPK 43123, UAS-RNAi Sec6 105836 and 22079, UAS-RNAi Sec8 45032 and 105653, UAS-RNAi Sec15 35162 and 35161, UAS-RNAi Vps34 107602, UAS-RNAi Atg8a 43096, UAS-RNAi Atg8b 17079, UAS-RNAi Rab11 108297, UAS-RNAi PVR 105353 were from VDRC (Dietzl et al., 2007). R3-*hml*Δ-*Gal4*, *UAS-2xeGFP* (Honti et al., 2013), UAS-RasV12, UAS-RasV12_S35, UAS-RasV12_G37, UAS-RasV12_C40 (Karim & Rubin, 1998), UAS-RalaWT, UAS-RalaG20V, UAS-RalaS25N (Sawamoto et al., 1999) and UAS-Rgl (Mirey et al., 2003), UAS-RNAi Vps15 (Abe et al., 2009), UAS-PVR (Duchek, Somogyi, Jékely, Beccari, & Rørth, 2001), UAS-Stat92EΔNΔC (Ekas et al., 2010) have been reported elsewhere. See a full list of stocks used in Table S2. The crosses were performed on German fly food (recipe available at http://flystocks.bio.indiana.edu/Fly_Work/media-recipes/germanfood.htm) supplemented with food coloring agent to allow visualization of the gut contents. The emptying of the gut marks the transition from wandering to resting third instar larvae, and only wandering third instar larvae were analyzed. Crosses were set up at 25 °C and shifted to 29 °C 66 h later to maximize Gal4 activity. Larvae were dissected under a UV lamp (Nightsea, GFP filter) for easy identification of GFP-positive lymph glands.

### Cloning and transgenics

pGEX-Sec5-RBD and pGEX-RalBP1-RBD encoding GST-tagged Ral-binding domains (RBDs) of human Sec5 and RalBP1 were a kind gift from A. Saltiel, University of Michigan. The sequences corresponding to the RBDs of *D. melanogaster* Sec5 and Rlip/RalBP1 were amplified from the plasmids SD03467 and GH01995 (DGRC), respectively, with primers 5’-ATAGAATTCGCGCCGCAGCCAGTGGTTAC-3’ and 5’-ATAGCGGCCGCCCGACCCCAGGCAAAATTCTG-3’ (Sec5) and 5’-ATAGAATTCGACATCCAGACGGAGTTGCG-3’ and 5’-ATAGCGGCCGCCTTGAGCCTATAGACTTCGTTG-3’ (Rlip/RalBP1) and inserted into EcoRI and NotI sites of pGEX-4T-1.

To generate pMet-3xFlag-RalaWT and –G20V plasmids, the Rala sequence was amplified from the plasmid LD21679 (DGRC) with primers 5’-ATAGGTACCatgGACTACAAAGACCATGACGGTGATTATAAAGATCATGACATCGATTACAAGGATGACGATGACAAGGaaAGCAAGAAGCCGACAGC-3’ (to add N-terminal3xFlag) or 5’-ATAggtaccATGAGCAAGAAGCCGACAGCCGGACCGGCGCTCCACAAGGTCATAATGGTGGGCAGTGTCGGCGTGGGAAAGTCC-3’ (to make the G20V mutation) and 5’-ATAGCGGCCGCATGAGCAAGAAGCCGACAGC-3’ and inserted into pMet using KpnI and NotI sites. All constructs were verified by sequencing.

The RalGPS (CG5522) coding sequence was amplified from the plasmid LD24677 (DGRC) with primers 5’-ATAGCGGCCGCATGATGCGATACTCGGAAATCTC-3’ and 5’-ATAGGTACCGCCCGGCTTATTCAAAGGACATTAGG-3’ and inserted into pUASTAttB using NotI and KpnI sites. Transgenics were generated by φC31-mediated site-specific integration (Bischof, Maeda, Hediger, Karch, & Basler, 2007) into attP154.

### Cell culture, lysates, Ral-GTP pulldown and western blotting

*Drosophila* S2 cells were cultured at 27 ºC in Schneider medium (Invitrogen) supplemented with 10% fetal bovine serum. S2 cells stably transfected with the following constructs have been described elsewhere: *pHS-SevS11* (Therrien et al., 1998), *pMet-EGFR* (a gift from N. Perrimon). pMet driven expression of EGFR was induced by adding 0.7 mM CuSO4 24 h prior to lysis and the cells were stimulated with the supernatant from Spitz secreting cells (Schweitzer, Shaharabany, Seger, & Shilo, 1995). *pHS-SevS11* cells were induced for 30 min at 37 ºC and incubated for the indicated time at 27 ºC before lysis. For insulin stimulation, S2 cells were treated with 20 μg/mL human recombinant insulin (Life Technologies #12585-014) for 0 to 80 min. For western blot analysis of S2 cell lysates, cells were lysed in ice-cold lysis buffer (20 mM Tris, pH 8.0, 137 mM NaCl, 10% glycerol, 1% NP-40, 1 mM EDTA) supplemented with 1X phosphatase inhibitor cocktail (Sigma #P2850), 10 μg/mL each aprotinin (Millipore Sigma #A6103) and leupeptin (Millipore Sigma #L2884), and 1 mM PMSF (Millipore Sigma #P7626). Lysates were then clarified by centrifugation at 10,000 x *g*.

Recombinant GST-tagged Ral-Binding Domains (RBDs) of human or *Drosophila* Sec5 or Rlip/RalBP1 were purified from bacteria by glutathione-Sepharose according to manufacturer’s instructions. S2 cells were lysed in cold RBD buffer (100 mM Tris pH 7.5, 150 mM NaCl, 10% glycerol, 5mM magnesium chloride, 1% NP-40, 1mM EDTA) supplemented with 1X phosphatase inhibitor cocktail (Sigma #P2850), 10 μg/mL each aprotinin and leupeptin, and 1 mM PMSF. 15 μg GST-fusion proteins immobilized on GSH-beads was added to lysates of equal protein concentration and incubated for 2 h at 4 ^o^C. After 3 washes in RBD buffer, the supernatant was completely removed, and bound proteins eluted by boiling in 2x Laemmli buffer.

For immunoblotting from fat body lysates, Rala transgenes or Rala RNAi was driven in the fat body by cg-Gal4. The fat bodies were dissected out and transferred to ice-cold lysis buffer (20 mM Tris, pH 8.0, 137 mM NaCl, 10% glycerol, 1% NP-40, 1 mM EDTA) supplemented with 1X phosphatase inhibitor cocktail (Sigma #P2850), 10 μg/mL each aprotinin (Millipore Sigma #A6103) and leupeptin (Millipore Sigma #L2884), and 1 mM PMSF (Millipore Sigma #P7626). Lysates were then clarified by centrifugation at 10,000 x g.

Protein samples were resolved by electrophoresis on 10-12% SDS-polyacrylamide gels and transferred to nitrocellulose membranes (Pall #66485). Specific proteins were detected using the following antibodies: anti-human RalB (1:1000; Proteintech 12340-1-AP), anti-GST (1:2000; Cell Signaling #2622), anti-MAPK (1:1000; Cell Signaling #4695); anti-AKT (1:2000; Cell Signaling #4691); anti-Actin (1:5000; Chemicon #MAB1501), anti-DER was kindly provided by G.M. Rubin, anti-pMAPK (1:1000; Cell Signaling #4370), anti-pAkt (1:1000; #4060), anti-sevenless was kindly provided by B.-Z. Shilo, anti-Flag (1:2000; Sigma F2555).

### Production of dsRNA and RNAi experiments in S2 cells

dsRNAs were generated as previously described (Clemens et al., 2000) with slight modifications. Sequences to target were selected from the DRSC database or designed when needed. DNA fragments (200-300 bp) containing coding sequences for the targeted proteins were amplified by PCR. Each PCR primer contained a 5’-T7 RNA polymerase binding site (GAATTAATACGACTCACTATAGGGAGA) followed by 21 nucleotides corresponding to the targeted sequence (see Table S3). 1 μg PCR product was used per T7 in vitro transcription reaction and incubated for 16 h. dsRNAs were generated by heating RNA samples to 95 °C and annealed by slow cooling to room temperature, followed by 30 min DNase I treatment. dsRNAs were NaOAc/EtOH-extracted, ethanol-precipitated, and resuspended in RNase-free H2O. dsRNA quality was verified on 2 % agarose gels.

For RNAi experiments, 10 × 10^6^ S2 cells were plated per 10 cm tissue culture dish (Nunc) with 10 μg/mL of the indicated dsRNAs and harvested 4 days later.

### Quantification of circulating hemocytes

To count circulating hemocytes, single third instar wandering larvae were bled in ESF921 (Expression systems) and transferred to individual wells of glass-bottom 384 well plates (Greiner μ-clear plate). The hemocytes were left to adhere for 1 h before fixation in 4% PFA, in PBS, followed by washes in PBS with 0.2% Triton X-100 (PBT 0.2%) and counterstaining of the nuclei with DAPI and the actin cytoskeleton with phalloidin-Alexa555 (Invitrogen). Mowiol was added to the wells before automated imaging with Operetta (PerkinElmer) or ScanR (Olympus) using a 20X objective. For each genotype, the number of GFP-positive hemocytes was quantified from at least 10 images from three individual larvae in four independent experiments using the Harmony (PerkinElmer) or ScanR analysis (Olympus) software. Identical analysis settings were applied for all samples within one experiment.

### Imaging of entire larvae

To image entire larvae, wandering larvae of indicated genotypes were collected, washed in PBS with 1% Triton X-100 and put to sleep into a chamber saturated with ethyl ether for 25 min. Anesthetized larvae were immobilized on a slide covered with double stick tape and immersed in glycerol before addition of a coverslip. Images were taken with an inverted microscope (Leica DM IRB) using a 2,5X objective. Alternatively, larvae were heat fixed. For this, larvae were washed as above, dried and put in a drop of glycerol on a microscope slide. The slide was placed on top of a heat block at 70 °C until the larvae stopped moving (this takes less than 10 s). A coverslip was added on top of the heat fixed larvae before imaging with a Leica MZFLIII stereomicroscope and LAS v4.9 Software.

### Immunofluorescence and microscopy

For immunofluorescence microscopy of lymph glands, larvae were dissected in ESF921 (Expression systems) supplemented with 1mM CaCl2, fixed for 15 min in 4% paraformaldehyde (PFA) in PBS on ice and washed three times in PBT 0.2%. The following conditions were used for each primary antibody: Hnt (1:10; DSHB 1G9, deposited by H.D. Lipshitz), Antp (1:100; DSHB 4C3, deposited by D. Brower), P1 (1:300; I. Ando (Kurucz, Markus, et al., 2007)), L1 (1:100; I Ando (Kurucz, Vaczi, et al., 2007)), Pvr (1:400; B. Shilo (Rosin, Schejter, Volk, & Shilo, 2004)), human RalB (1:100; Proteintech 12340-1-AP). Lymph glands were incubated overnight at 4 °C with primary antibodies, washed three times in PBT 0,2% and incubated at room temperature for 2 h with fluorophore-conjugated secondary antibodies (1:500) from Molecular Probes. Lymph glands were counterstained with 100 ng/mL DAPI or Hoechst, washed twice in PBT 0,2% and mounted in Mowiol (Sigma). Confocal imaging was performed with three different confocal microscopes. First a LSM510 (Carl Zeiss) confocal microscope, equipped with an Ar-laser multiline (458/488/514 nm), a laser diode 405–30 CW (405 nm), and two HeNe lasers (543 and 633 nm). The objective used with LSM510 was a Plan-Neofluar 40x/1.3 Oil DIC (Carl Zeiss). Alternatively, a Leica TCS SP8 confocal microscope equipped with a Plan-Apochromat 40x/1.3 Oil DIC objective, a UV (405nm) laser and a continuous wavelength (CW) white light laser set to 488 nm and 594 nm was used. Finally, a Zeiss LSM880 equipped with an Ar-laser multiline (458/488/514 nm), a DPSS-561 10 (561 nm), a laser diode 405–30 CW (405 nm), and an HeNe laser (633 nm). The objective used was a Plan-Apochromat 63x/1.4 Oil DIC III (Carl Zeiss).

The overview images of entire lymph glands were acquired by a Zeiss Axio-imager using a 10X objective.

For immunofluorescence imaging of circulating hemocytes, third instar wandering larvae were gently opened in PBS and the solution was transferred to individual wells of concanavalin A-coated 15 well microscope slides (MP Biomedicals) in triplicates. The hemocytes were left to adhere for 2 h before fixation in 4% PFA in PBS, followed by washes in PBT 0.2% and o.n. incubation with indicated antibodies. Hemocytes were then stained with secondary antibodies, counterstained with DAPI and imaged by confocal microscopy as described for lymph glands.

### Image processing and quantification

For direct intensity comparisons, images were captured with identical settings below pixel value saturation and post-processed identically. Microscopy images were processed in ImageJ or Adobe Photoshop. Brightness and contrast were adjusted. Maximal intensity projections were created in ImageJ or Zen 2012 software (Zeiss). Images were cropped to show an entire lymph gland or a region of interest.

Quantification of Hnt+ compared to total DAPI+ cells was performed with the Plot applet of Imaris software (Imaris 7.7.2, Bitplane AG, Zürich, Switzerland). Identical analysis settings were applied for all samples within one experiment.

### Quantification of lymph gland size by flow cytometry analysis

To quantify the lymph gland size, 15 glands were dissected and put immediately in 100 μl of 1X trypsin-EDTA (No phenol red; Life technologies) diluted in PBS, incubated 15 min at 25 °C and pipet up and down 40 times through a 200 μl siliconized tip. Trypsin was neutralized by adding 300 ul of 2% FBS in PBS and the cell suspension was filtered using a FACS tube with a cell-strainer cap (Falcon). The percentage of GFP+ cells was determined by flow cytometry (BD FACSCanto II or LSRII). The total number of cells per sample was quantified by counting cells with a hemocytometer. The number of GFP+ cells per lymph gland was determined the following way: %GFP+ cells x Total number of cell per sample/15 lymph glands.

### qPCR analysis

For qPCR analysis, 15 lymph glands were dissected and put immediately in RPL buffer (RNeasy Micro kit) and RNA was extracted following the standard procedure of the kit. Reverse transcription was performed on 250 μg of RNA using the High capacity reverse transcription kit from Applied Biosystems. Taqman qPCR assays were designed using the Universal Probe Library design center (Roche). SYBRgreen qPCR primers were selected from http://www.flyrnai.org/flyprimerbank (Hu et al., 2013). The primer sequences and the Universal ProbeLibrary probe number are listed in Table S4. Assays were designed such that the amplified regions did not overlap with sequences targeted by dsRNA. Taqman qPCR reactions were performed with the TaqMan® Real-Time PCR Master Mix and analyzed with the ViiA™ 7 Real-Time PCR System. SYBRgreen qPCR reactions were performed with the Fast SYBR Green Master Mix (Applied Biosystems) and analyzed with the Applied Biosystems StepOnePlus Real-time PCR system. The quantification of target genes was determined using the Ct method. Briefly, the Ct (threshold cycle) values of target genes were normalized to a reference gene (Act5C unless indicated otherwise) where ΔCt = Ct_target_ – Ct_Act5C_, and then compared with a calibrator sample (RLuc RNAi or WT) where ΔΔCt = ΔCt_Sample_ - ΔCt_Calibrator_. Relative expression (RQ) was calculated with the formula RQ = 2^-ΔΔCT^. The RNAi efficiencies measured for all lines tested are listed in Table S5.

### Transcriptome analysis

RNA-seq libraries were prepared from 200 ng of RNA using the KAPA stranded mRNA-seq Kit (KAPABiosystems). Sequencing was performed on an Illumina HiSeq2000 instrument using TruSeq SBS v3 chemistry. Sequences were trimmed for sequencing adapters and then aligned to the reference *D. melanogaster* BDGP6 genome using the STAR software (version 2.7.1a). Expression values were estimated for genes and transcripts defined in Ensembl (release 99) using the RSEM algorithm (version 1.2.28) and then normalized across samples as TPM values. The sequenced data from these experiments are available at Gene Expression Omnibus accession: GSE148035.

### Statistics

Statistical analysis was performed using Graphpad Prism. The data was assumed to be normally distributed. For multiple comparisons, one-way ANOVA with Bonferroni post-testing was used. Data shown as normalized to WT/control has been normalized to the mean of all the control values from the different experiments. The exception is qPCR experiments to test for target knockdown, where target expression was set to 1 in each experiment.

## Supporting information

Supplementary methods and references

## Acknowledgements

We would like to thank all the researchers who generously shared reagents: H. Okano, M. Balakireva, S. Goto, G.M. Rubin, A. Saltiel, N. Perrimon, P. Rorth, I. Ando, B. Shilo and E.Bach. Stocks obtained from the Bloomington Drosophila Stock Center (NIH P40OD018537) and Vienna Drosophila Resource Center (VDRC) were used in this study. Several antibodies were obtained from the Developmental Studies Hybridoma Bank created by the NICHD of the NIH and maintained at The University of Iowa. Several plasmids were obtained from Drosophila Genomics Resource Center, supported by NIH grant 2P40OD010949. We thank K.M. Dahlgren and F. Lussier for technical assistance. We thank Dr. Ellen Tenstad/Science Shaped for designing the graphical figures. The Bioimaging, Bioinformatics, Flow cytometry, and Genomics core facilities of IRIC as well as the core facilities for Advanced Light Microscopy and Flow Cytometry at Oslo University Hospital, Gaustad and Montebello nodes, are acknowledged for access, help and services. H.K. was supported by fellowship 221552/F20 from the Research Council of Norway and fellowship 2017062 from the South-Eastern Norway Regional Health Authority. J.M.E. was supported by Norwegian Cancer Society projects 4487303, 182524 and 208012, and Norwegian Research Council projects 261936 and 294916. M.T. is recipient of Tier 1 Canada Research Chair in Intracellular Signaling. This work was supported by the Research Council of Norway through its Centres of Excellence funding scheme, project number 262652. This project was also supported by funds from the Canadian Institutes for Health Research (MOP-15375 and FDN 388023) to M.T.

## Competing interests

The authors declare that no competing interests exist.

## Notes

https://www.ncbi.nlm.nih.gov/geo/query/acc.cgi?acc=GSE148035

### Competing Interest Statement

The authors have declared no competing interest.

