## Supplementary methods and references for "Rala and the exocyst control Pvr trafficking and signaling to ensure lymph gland homeostasis in *Drosophila melanogaster*"

*Quantification of phagocytic activity*

To evaluate phagocytic activity of larval circulating hemocytes, 15 larvae were washed in PBT 1%, transferred to PBS 1× and then dried on a paper towel. Cleaned larvae were bled in 110 µl of ESF921 medium and circulating hemocytes were counted using a hemocytometer. Equivalent numbers of hemocytes per sample and per genotype were incubated with 1 µl, 2.5 µl and 5 µl of Alexa-488 labeled E. Coli bioparticles (Life Technologies) for 20 min at room temperature. Fluorescence of extracellular bioparticles was quenched with 50 µl of 4% Trypan blue and thereafter the samples were immediately put on ice. Before analysis by flow cytometry (BD FACSCanto II), 300 µl of PBS 1× +2% FBS was added and samples were filtered into FACS tubes through a cell-strainer cap. The mean fluorescence intensity per cell was used as a parameter to calculate phagocytic activity. This protocol was adapted from (Neyen et al., 2014). For bioparticle preparation, 2 mg of bioparticle powder were vortexed in 2 ml of PBS 1× for 1 min and then passed 40 times through a 26-gauge needle. 20 µl aliquots were stored at −80 °C for later use.
