## Supplementary table 1 for "Rala and the exocyst control Pvr trafficking and signaling to ensure lymph gland homeostasis in *Drosophila melanogaster*"

**Table S1: mRNA levels of Rala-related proteins in entire lymph glands**

| Gene name | CG number | Average (TPM) | SD |
|---|---|---|---|
| Act5C | CG4027 | 2997,83 | 1019,04 |
| Atg1 | CG10967 | 16,50 | 1,83 |
| Atg12 | CG10861 | 19,65 | 3,10 |
| Atg16 | CG31033 | 14,47 | 1,00 |
| Atg18a | CG7986 | 48,40 | 2,74 |
| Atg18b | CG8678 | 6,99 | 0,86 |
| Atg4a | CG4428 | 36,77 | 4,20 |
| Atg4b | CG6194 | 8,98 | 1,46 |
| Atg5 | CG1643 | 20,90 | 1,47 |
| Atg6 | CG5429 | 21,10 | 1,56 |
| Atg7 | CG5489 | 13,29 | 3,54 |
| Atg8a | CG32672 | 330,87 | 28,99 |
| Atg8b | CG12334 | 0,00 | 0,00 |
| Exo70 | CG7127 | 20,05 | 0,98 |
| Exo84 | CG6095 | 20,19 | 0,84 |
| Gapdh1 | CG12055 | 336,53 | 35,30 |
| RalGAPα | CG5521 | 7,37 | 1,03 |
| RalGAPβ | CG34408 | 16,91 | 1,35 |
| RalGPS | CG5522 | 18,78 | 1,99 |
| Pi3K59F | CG5373 | 11,56 | 0,57 |
| Pvr | CG8222 | 135,20 | 13,00 |
| Rab11 | CG5771 | 449,53 | 67,77 |
| Rala | CG2849 | 40,86 | 4,77 |
| Ras85D | CG9375 | 94,13 | 9,72 |
| Rgl | CG8865 | 4,88 | 0,91 |
| Rlip/RalBP1 | CG11622 | 15,14 | 0,98 |
| rl/MAPK | CG12559 | 65,61 | 2,02 |
| RtGEF | CG10043 | 15,46 | 0,94 |
| Sec10 | CG6159 | 25,31 | 3,01 |
| Sec15 | CG7034 | 13,56 | 1,23 |
| Sec3 | CG3885 | 17,10 | 1,66 |
| Sec5 | CG8843 | 16,97 | 0,92 |
| Sec6 | CG5341 | 15,00 | 0,53 |
| Sec8 | CG2095 | 41,70 | 1,65 |
| Vps15 | CG9746 | 8,50 | 0,54 |
