## Supplementary table 2 for "Rala and the exocyst control Pvr trafficking and signaling to ensure lymph gland homeostasis in *Drosophila melanogaster*"

**Table S2: Fly stocks used**

| Name | Obtained from | Stock number | Comment/Reference |
|---|---|---|---|
| *hmlΔ-Gal4, UAS-2XeGFP* | BDSC | BL30140 | |
| *hmlΔ-Gal4, UAS-2XeGFP* | BDSC | BL30142 | |
| R3-*hmlΔ-Gal4*, *UAS-2xeGFP* | | | (Honti et al., 2013) |
| P{w[+mC]=Cg-GAL4.A, UAS-GFP | BDSC | BL7011 | Recombined with UAS-GFP |
| UAS-RNAi Rala #1 | VDRC | 43622 | |
| UAS-RNAi Rala #2 | VDRC | 105296 | |
| UAS-RNAi Rala #3 | BDSC | BL28950 | |
| UAS-RNAi Rala #4 | BDSC | BL34375 | |
| UAS-RNAi Sec5 #1 | VDRC | 28873 | |
| UAS-RNAi Sec5 #2 | VDRC | 28874 | |
| UAS-RNAi Sec5 #3 | BDSC | BL27526 | |
| UAS-RNAi Rlip #1 | VDRC | 101635 | |
| UAS-RNAi Rlip #2 | VDRC | 16244 | |
| UAS-RNAi Exo84 #1 | VDRC | 108650 | |
| UAS-RNAi Exo84 #2 | BDSC | BL28712 | |
| UAS-RNAi Exo84 #3 | VDRC | 30112 | |
| UAS-RNAi Sec6 #1 | VDRC | 105836 | |
| UAS-RNAi Sec6 #2 | VDRC | 22079 | |
| UAS-RNAi Sec8 #1 | VDRC | 45032 | |
| UAS-RNAi Sec8 #2 | VDRC | 105653 | |
| UAS-RNAi Sec15 #1 | VDRC | 35162 | |
| UAS-RNAi Sec15 #2 | VDRC | 35161 | |
| UAS-RNAi RalGPS/CG5522 #1 | VDRC | 40595 | |
| UAS-RNAi RalGPS/CG5522 #2 | VDRC | 40596 | |
| UAS-RasV12 on 2nd | | | (Karim and Rubin, 1998) |
| UAS-RasV12 on 3rd | | | (Karim and Rubin, 1998) |
| UAS-RasV12_S35 | | | (Karim and Rubin, 1998) |
| UAS-RasV12_G37 on 2nd | | | (Karim and Rubin, 1998) |
| UAS-RasV12_G37 on X | | | (Karim and Rubin, 1998) |
| UAS-RasV12_C40 | | | (Karim and Rubin, 1998) |
| UAS-RalaWT | | | (Sawamoto et al., 1999) |
| UAS-RalaG20V | | | (Sawamoto et al., 1999) |
| UAS-RalaS25N | | | (Sawamoto et al., 1999) |
| UAS-Rgl | | | (Mirey et al., 2003) |
| UAS-RalGPS | | | This study |
| UAS-RNAi mapk #1 | BDSC | BL36058 | |
| UAS-RNAi mapk #2 | VDRC | 43123 | |
| UAS-RNAi mapk #3 | BDSC | BL34855 | |
| UAS-RNAi Atg1 | BDSC | BL26731 | |
| UAS-RNAi Atg8a | VDRC | 43096 | |
| UAS-RNAi Atg8b | VDRC | 17079 | |
| UAS-RNAi Atg7 | BDSC | BL27707 | |
| UAS-RNAi Atg5 | BDSC | BL27551 | |
| UAS-RNAi Atg12 | BDSC | BL27552 | |
| UAS-RNAi Atg4 | BDSC | BL35740 | |
| UAS-RNAi Atg16 | BDSC | BL28060 | |
| UAS-RNAi Vps34 | VDRC | 107602 | |

| | | | |
|---|---|---|---|
| UAS-RNAi Vps15 | | | (Abe et al., 2009) |
| UAS-RNAi Atg6 | BDSC | BL35741 | |
| UAS-RNAi Atg18 | BDSC | BL34714 | |
| UAS-RNAi Rab11 #1 | BDSC | BL27730 | |
| UAS-RNAi Rab11 #2 | VDRC | 108297 | |
| UAS-PVR | | | (Duchek et al., 2001) |
| UAS-Stat92EΔNΔC | | | (Ekas et al., 2010) |
| UAS-RNAi PVR | VDRC | 105353 | |
