## Supplementary table 3 for "Rala and the exocyst control Pvr trafficking and signaling to ensure lymph gland homeostasis in *Drosophila melanogaster*"

**Table S3: dsRNAs for RNAi in S2 cell culture**

| Target | 5'primer | 3'primer | Reference |
|---|---|---|---|
| RLuc (control) | TGCTGGACAGCTTCATCAAC | ATGTCCTCCTCGATGTCTGG | our design |
| Rala | CTTTCGTCGTCTGTGATGGA | GCGCTCCACAAGGTCATAAT | DRSC27866 |
| RalGPS (CG5522) #1 | CCAATGGACACACTTTGCAC | ATGATTCGGTCTTGGAGCAC | DRSC27139 |
| RalGPS (CG5522) #2 | GTCCAAAAGCTCGTGTGGTT | GCAAGAATTCCATCGGTTGT | DRSC37170 |
| Rgl #1 | CATAATAGAAGCCGTCGATG | CGGAGCTGGCGAATATTT | our design |
| Rgl #2 | AATGGTCATTCGTGGCTTTC | CAGGATACTGGACACCACCC | our design |
| Rgl RA | CAGGCAGACGCATAGACGTA | GAGATTTCCACGGATGCACT | our design |
| RalGAPα (CG5521) #1 | TTTGTCGGCCAGCTCTAACT | CTTCGATGTCAGTTGCTCCA | DRSC30087 |
| RalGAPα (CG5521) #2 | GGAGCATTCAGCTGGCTTAC | CTTGCTGGGACGCTTCTATC | DRSC36444 |
| RalGAPβ (CG34408) | CTGCAGCAGTTCATCCACTC | GGTGACCCATTATGGCATTC | DRSC28674 |
| TD-60 (CG9135) #1 | TCGGTTTCGGGCGTAATC | ATTATTGCCGCAAGCAAAG | DRSC03144 |
| TD-60 (CG9135) #2 | TTTTCGATACTCAGGGTCGG | CAACACCGTGTGCGAGTAAC | our design |
