## Supplementary table 4 for "Rala and the exocyst control Pvr trafficking and signaling to ensure lymph gland homeostasis in *Drosophila melanogaster*"

**Table S4: qPCR assays**

| Target | Primer 1 | Primer 2 | Taqman probe # |
|---|---|---|---|
| Rala | gcaacccaggaattcagaga | tcattcagatcgcacttgttg | 18 |
| Rala | gactacgagcccaccaagg | tgcgaaagtagttatctctgatgg | 79 |
| RalGPS (CG5522) | ccacggataatctgcgctac | tgctgctgctgttgtgaag | 63 |
| Rgl | cctgaagaaagtgcgatacca | tgggaaatctcatcatctgagtc | 76 |
| Rgl-RA | gacaagaagcgcaaggagtt | tccgcctggagattatatgtg | 76 |
| RalGAPα (CG5521) | ccgaggcgaatctaaaacaa | gcagaaggcacaggatcttc | 46 |
| RalGAPβ (CG34408) | tgccaagtttaagtactttgtgacc | ggaccgcgtatgagcaga | 22 |
| TD-60 (CG9135) | ctttgcttgcggcaataatc | atttcgtccttgggctctg | 45 |
| Exo84 | cgacgattttaattttagcgttg | tgtaggcctcgatttcctttt | 87 |
| Sec5 | cgtcaatgagactgccaaga | acgcatgttccgcttgac | 50 |
| Rlip | cagatgtctcaccgactaatgg | gatgttaaagggggcacatactt | 53 |
| Act5c | accgagcgcggttactct | cttgatgtcacggacgatttc | 77 |
| Atg1 | CGTCAGCCTGGTCATGGAGTA | TAACGGTATCCTCGCTGAG | SYBRgreen |
| Atg7 | CGACTCGAAACGCTCCATTA | ATTCTGTTGTAGGCCGTGTAG | SYBRgreen |
| Vps34 | CTCCATCTCAATGTTCGGCAA | GCCGTTCCTGTGTTTCTTGG | SYBRgreen |
| Rab11 | ATTTGCTCTCACGTTTCACGC | GCCATCGACCTCTATGCTGC | SYBRgreen |
| Ras85D | TGCGGGATCAGTATATGCGGA | GCATCCTTTACGCGCTTGA | SYBRgreen |
| Gapdh1 | TAAATTCGACTCGACTCACGGT | CTCCACCACATACTCGGCTC | SYBRgreen |
