## Supplementary table 5 for "Rala and the exocyst control Pvr trafficking and signaling to ensure lymph gland homeostasis in *Drosophila melanogaster*"

**Table S5: RNAi efficiency measured as relative mRNA levels in entire lymph glands for all lines tested in this study**

| Hml-Gal4> | Stock number (VDRC or BDSC) | mRNA | housekeeping reference | Mean relative expression (compared to WT) | Relative expression standard deviation |
|---|---|---|---|---|---|
| UAS-RNAi Rala #1 | 43622 | Rala | rtgef | 0.67 | 0.35 |
| UAS-RNAi Rala #2 | 105296 | Rala | rtgef | 0.66 | 0.23 |
| UAS-RNAi Rala #3 | BL28950 | Rala | rtgef | 0.74 | 0.06 |
| UAS-RNAi Rala #4 | BL34375 | Rala | rtgef | 0.45 | 0.05 |
| UAS-RNAi Rgl | 23639 | Rgl | rtgef | 0.77 | 0.10 |
| UAS-RNAi Rgl | 106468 | Rgl | rtgef | 0.60 | 0.22 |
| UAS-RNAi Rgl | BL33389 | Rgl | rtgef | 1.12 | 0.31 |
| UAS-RNAi RalGPS/CG5522 #1 | 40595 | Rgl | rtgef | 1.16 | 0.21 |
| UAS-RNAi RalGPS/CG5522 #2 | 40596 | Rgl | rtgef | 1.22 | 0.03 |
| UAS-RNAi RalGPS/CG5522 | BL31142 | Rgl | rtgef | 0.56 | 0.16 |
| UAS-RNAi Rgl | 23639 | RalGPS/CG5522 | Act5c | 1.27 | 0.11 |
| UAS-RNAi Rgl | 106468 | RalGPS/CG5522 | Act5c | 0.18 | 0.03 |
| UAS-RNAi Rgl | BL33389 | RalGPS/CG5522 | Act5c | 0.44 | 0.03 |
| UAS-RNAi RalGPS/CG5522 #1 | 40595 | RalGPS/CG5522 | Act5c | 0.36 | 0.05 |
| UAS-RNAi RalGPS/CG5522 #2 | 40596 | RalGPS/CG5522 | Act5c | 0.32 | 0.07 |
| UAS-RNAi RalGPS/CG5522 | BL31142 | RalGPS/CG5522 | Act5c | 0.14 | 0.03 |
| UAS-RNAi Exo84 #1 | 108650 | Exo84 | Act5c | 0.71 | 0.10 |
| UAS-RNAi Exo84 #2 | BL28712 | Exo84 | Act5c | 0.47 | 0.10 |
| UAS-RNAi Exo84 #3 | 30112 | Exo84 | Act5c | 0.67 | 0.25 |
| UAS-RNAi Sec5 #1 | 28873 | Exo84 | Act5c | 1.35 | 0.51 |
| UAS-RNAi Sec5 #2 | 28874 | Exo84 | Act5c | 1.23 | 0.40 |
| UAS-RNAi Sec5 #3 | BL27526 | Exo84 | Act5c | 1.05 | 0.52 |
| UAS-RNAi Exo84 #1 | 108650 | Sec5 | Act5c | 1.33 | 0.20 |
| UAS-RNAi Exo84 #2 | BL28712 | Sec5 | Act5c | 0.88 | 0.10 |
| UAS-RNAi Exo84 #3 | 30112 | Sec5 | Act5c | 1.25 | 0.11 |
| UAS-RNAi Sec5 #1 | 28873 | Sec5 | Act5c | 0.68 | 0.14 |
| UAS-RNAi Sec5 #2 | 28874 | Sec5 | Act5c | 1.13 | 0.13 |

| | | | | | |
|---|---|---|---|---|---|
| UAS-RNAi Sec5 #3 | BL27526 | Sec5 | Act5c | 0.60 | 0.11 |
| UAS-RNAi RalGAPα | 22150 | RalGAPα | rtgef | 0.86 | 0.04 |
| UAS-RNAi RalGAPα | 110722 | RalGAPα | rtgef | 1.14 | 0.12 |
| UAS-RNAi RalGAPβ | 106188 | RalGAPα | rtgef | 1.18 | 0.17 |
| UAS-RNAi RalGAPβ | 106659 | RalGAPα | rtgef | 1.01 | 0.05 |
| UAS-RNAi RalGAPα | 22150 | RalGAPβ | rtgef | 1.17 | 0.13 |
| UAS-RNAi RalGAPα | 110722 | RalGAPβ | rtgef | 1.06 | 0.04 |
| UAS-RNAi RalGAPβ | 106188 | RalGAPβ | rtgef | 0.71 | 0.03 |
| UAS-RNAi RalGAPβ | 106659 | RalGAPβ | rtgef | 1.13 | 0.05 |
| UAS-RNAi Rlip #1 | 101635 | Rlip | Act5c | 1.02 | 0.24 |
| UAS-RNAi Rlip #2 | 16244 | Rlip | Act5c | 0.61 | 0.09 |
| UAS-RNAi Atg1 | BL26731 | Atg1 | gapdh | 0.53 | 0.18 |
| UAS-RNAi Atg7 | BL27707 | Atg7 | gapdh | 0.39 | 0.31 |
| UAS-RNAi Vps34 | 107602 | Vps34 | gapdh | 0.66 | 0.22 |
| UAS-RNAi Rab11 | BL27730 | Rab11 | gapdh | 0.60 | 0.07 |
| UAS-RNAi Exo70 #1 | 27867 | Exo70 | gapdh | 1.13 | 0.20 |
| UAS-RNAi Exo70 #2 | 103717 | Exo70 | gapdh | 0.93 | 0.17 |
| UAS-RNAi Sec10 | BL27483 | Sec10 | gapdh | 0.61 | 0.22 |
