## Supplementary figure 1 for "Rala and the exocyst control Pvr trafficking and signaling to ensure lymph gland homeostasis in *Drosophila melanogaster*"

**Supplementary figures:**

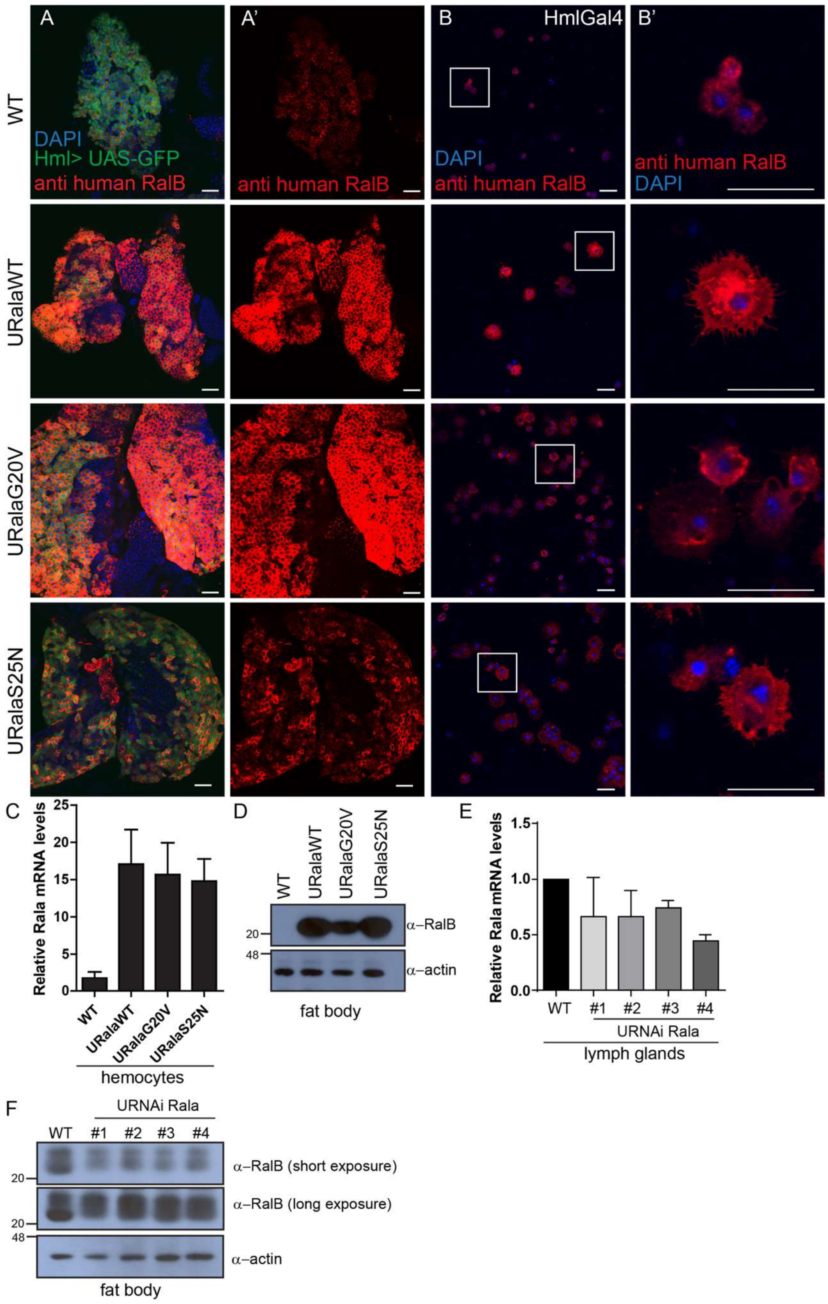


**Supplementary Figure 1. Detection of Rala overexpression in hemocytes**

**A-B.** Ectopic expression of *Dm* Rala in larval lymph glands (A-A', single confocal plane close to lymph gland surface) and circulating hemocytes (B-B', single confocal plane) was monitored by immunostaining with an antibody against human RalB. To be able to detect some Rala signal in WT lymph glands (A-A'), the signal upon Rala overexpression imaged at the same settings is partly saturated. Note that in both tissues, overexpressed, but not endogenous *Dm* Rala could be detected. **C,E.** Rala mRNA levels were measured from hemocytes or entire lymph glands by qRT-PCR. Graphs show mean normalized to the control (WT) +/- stdev from 3 independent experiments. **D,F.** Endogenous Rala was detected by immunoblotting of fat body lysate.
