## Supplementary figure 2 for "Rala and the exocyst control Pvr trafficking and signaling to ensure lymph gland homeostasis in *Drosophila melanogaster*"

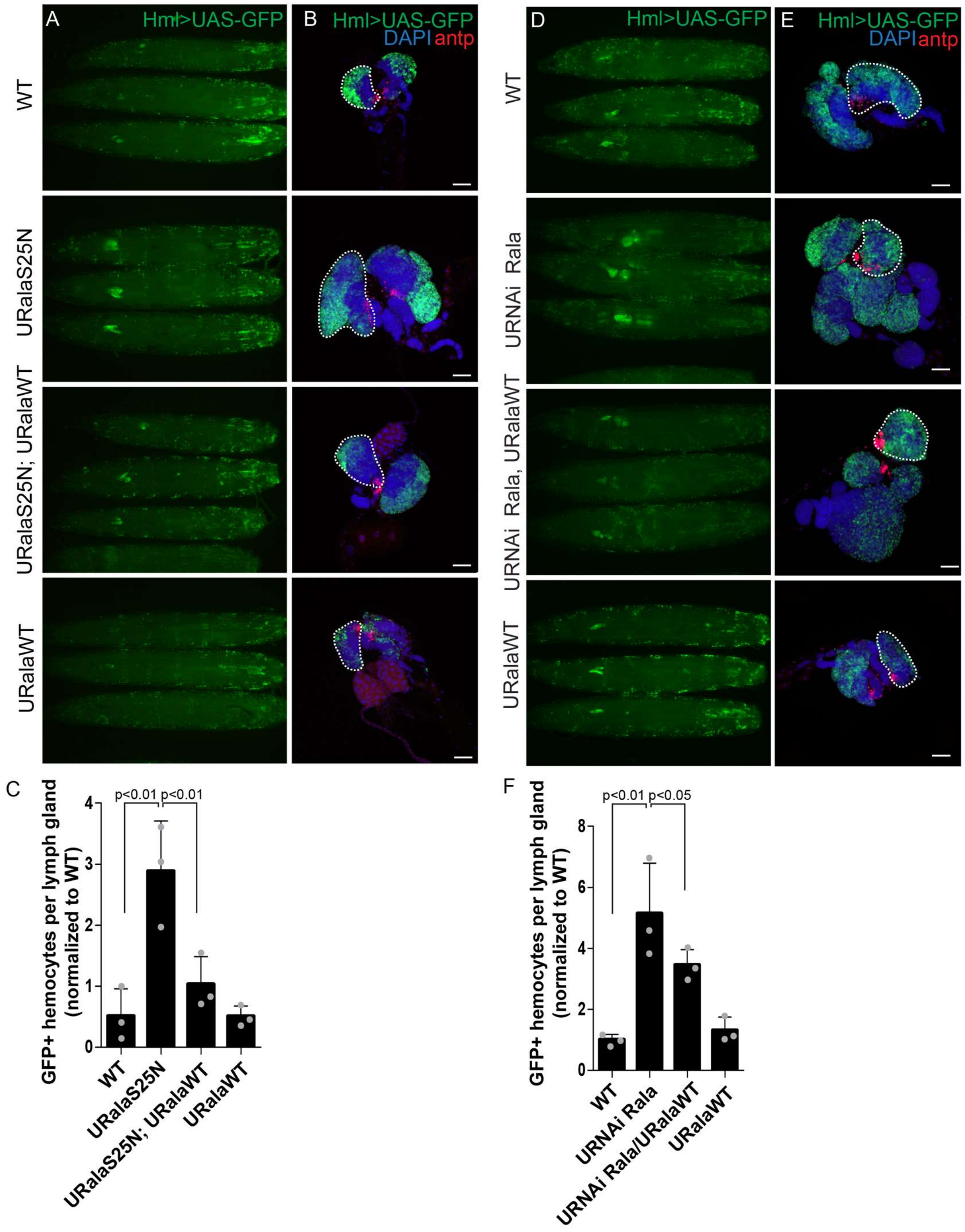


**Supplementary Figure 2. Rala RNAi and RalaS25N-induced lymph gland enlargement is rescued by RalaWT coexpression.**

**A and D.** Wandering third instar larvae expressing the indicated transgenes driven by Hml-Gal4 were imaged by widefield fluorescence microscopy. **B and E.** Representative widefield fluorescence images of lymph glands where the cortical zone is marked by Hml-driven GFP and the PSC by antp immunofluorescence. One primary lobe is outlined in each image. Scale bar 50 µm. **C and F.** Quantification of the number of GFP positive hemocytes per larval lymph gland. Graph shows average GFP-positive hemocytes per lymph gland normalized to control (WT) +/- stdev from 3 independent experiments. Statistically significant differences are indicated (one-way ANOVA).
