## Supplementary figure 3 for "Rala and the exocyst control Pvr trafficking and signaling to ensure lymph gland homeostasis in *Drosophila melanogaster*"

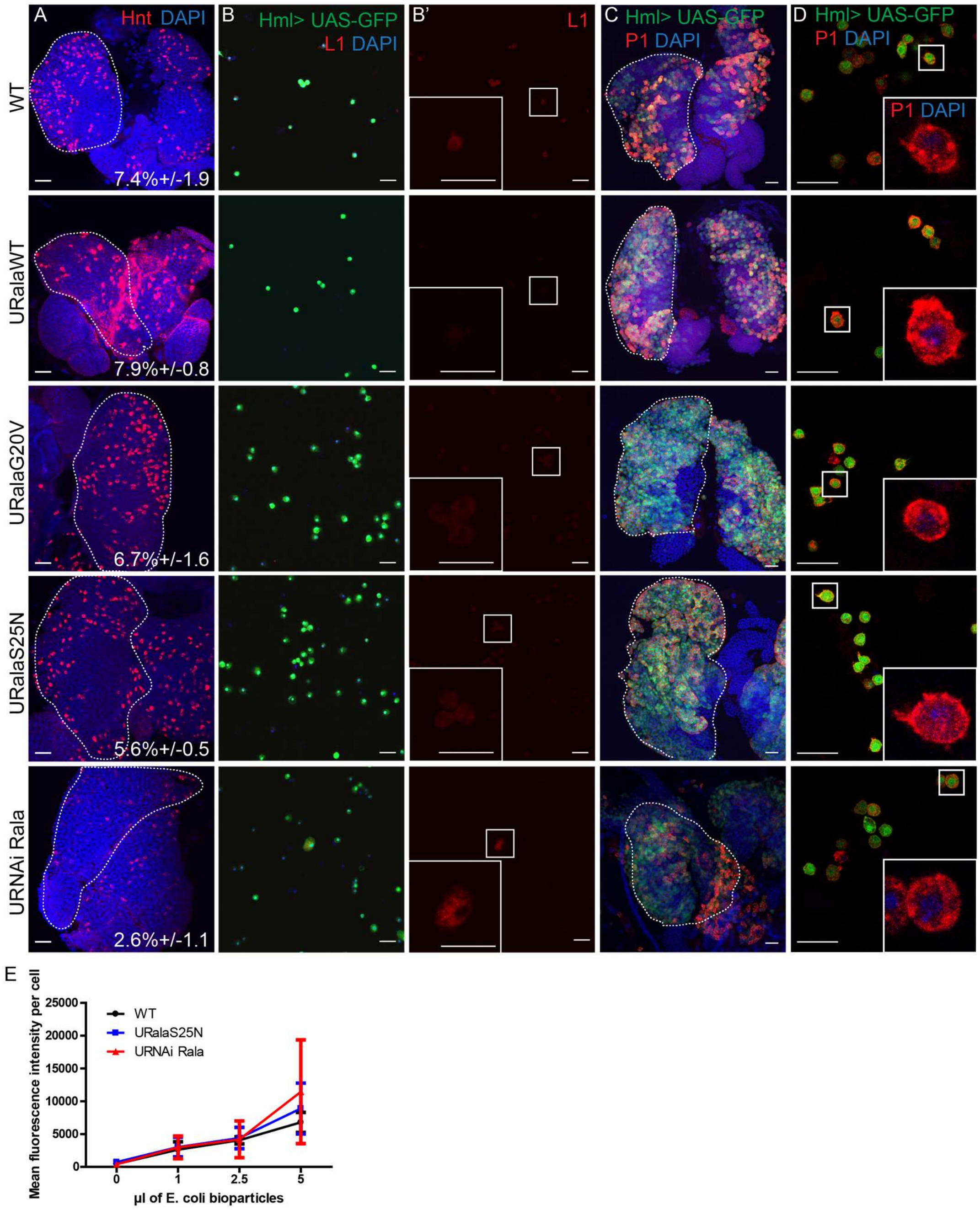


**Supplementary Figure 3**. **Effects of perturbed Rala activation on the differentiation of hemocytes.**

**A.** Representative maximum intensity projections of confocal stacks of lymph glands from the indicated genotypes stained with the crystal cell marker Hnt. The percentage of crystal cells per primary lobe was quantified and shown as mean +/- stdev, n=3. **B-B'.** Representative confocal images of circulating hemocytes immunostained with the lamellocyte marker L1. Note that lamellocytes are sometimes observed in Rala RNAi. **C-D.** Immunostaining of NimrodC1/P1 in lymph glands (C, maximum intensity projections of confocal stacks) and circulating hemocytes (D, single confocal planes) from the indicated genotypes stained with the plasmatocyte marker. The *R3-hmlΔ-Gal4, UAS-2xeGFP* line was used for panel C and D as it was demonstrated to be homozygous wild-type for the NimrodC1 gene (Honti et al., 2013). Scale bars 20 µm. **E.** WT, RalaS25N-expressing or Rala depleted hemocytes were challenged with fluorophore-labelled E. coli bioparticles. Phagocytosed bioparticles were quantified by flow cytometry.
