## Supplementary figure 4 for "Rala and the exocyst control Pvr trafficking and signaling to ensure lymph gland homeostasis in *Drosophila melanogaster*"

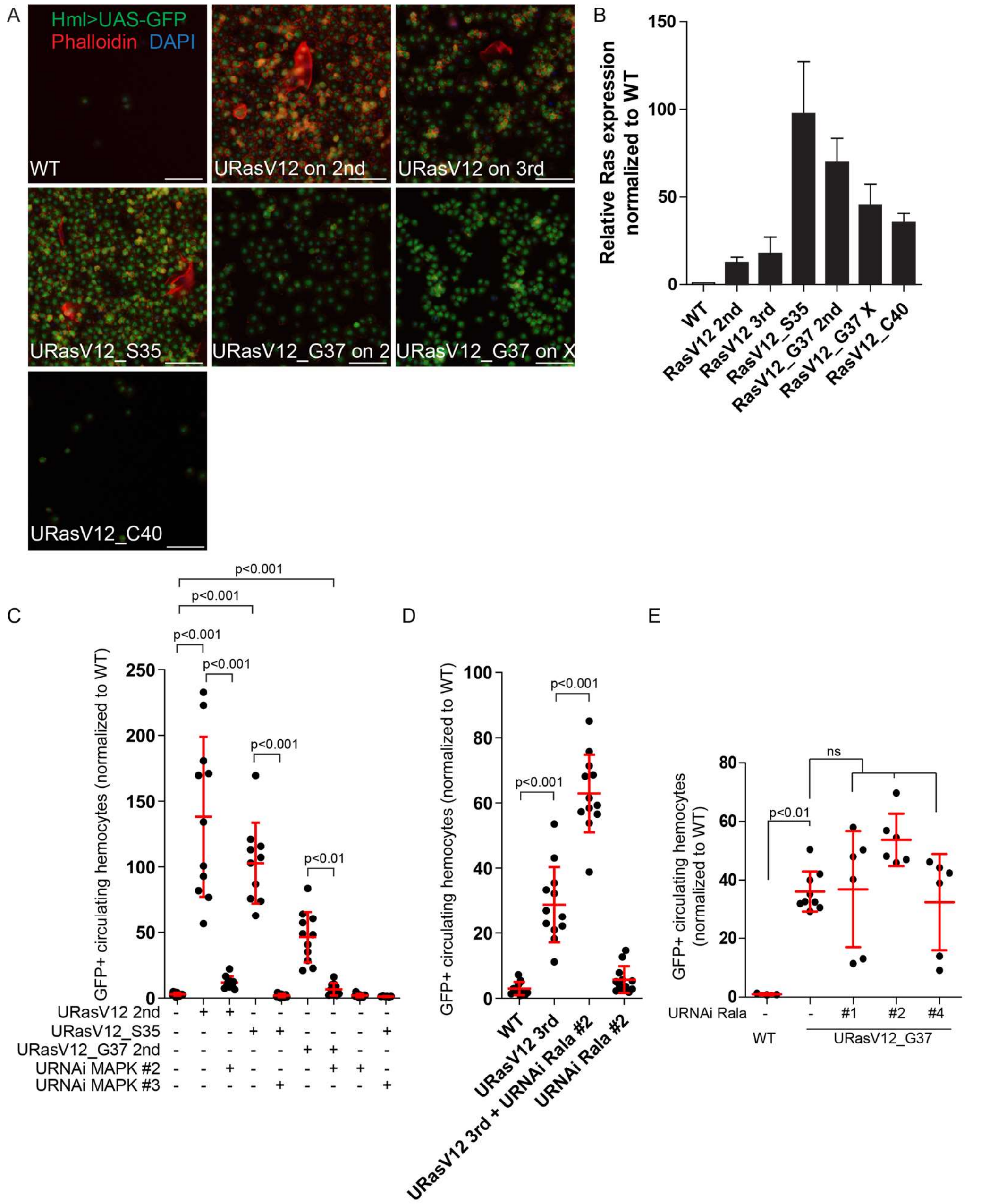


**Supplementary Figure 4. The RasV12G37-induced lymph gland overproliferation is not dependent on Rala.**

**A.** Widefield fluorescence images of hemolymph from larvae expressing the indicated transgenes in the hematopoietic system. Note the presence of enlarged actin-rich (Phalloidin-positive) hemocytes, that is lamellocytes, upon expression of RasV12 or RasV12_S35. Scale bars 50 µm. **B.** Quantification of Ras expression levels in circulating hemocytes by qPCR. The graph shows mean +/- stdev from 3 independent experiments. **C-E.** The number of GFP-positive circulating hemocytes from larvae expressing the indicated transgenes driven by Hml-Gal4 was quantified by high-content microscopy. The graph shows mean +/- stdev from a total of 12 larvae from 4 independent experiments (C-D) or from a total of 6 larvae from 3 independent experiments (E). Statistically significant differences are indicated (one-way ANOVA).
