## Supplementary figure 5 for "Rala and the exocyst control Pvr trafficking and signaling to ensure lymph gland homeostasis in *Drosophila melanogaster*"

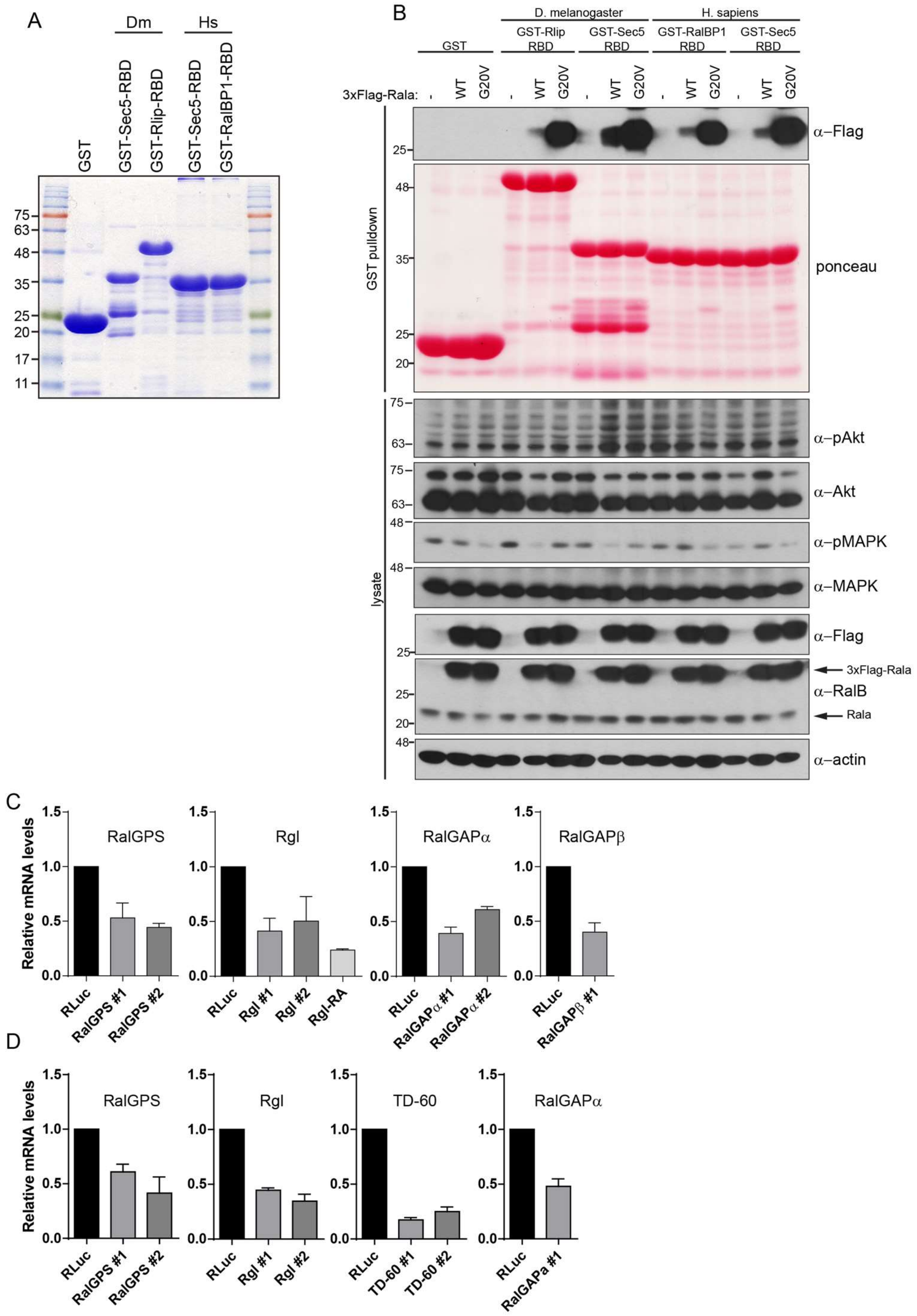


**Supplementary Figure 5. Characterization of Rala activity assay by GST pulldown.**

**A.** Coomassie blue stained SDS-PAGE gel showing the purified GST proteins. **B.** S2 cells were transiently transfected to express 3x-Flag-RalaWT or GTP-locked 3xFlag-RalaG20V. Recombinant GST-tagged Ral-binding domain (RBD) from human or fly Rlip/RalBP1 or Sec5 bound to GSH sepharose was incubated with the indicated lysates and the resulting pull-downs were analyzed by immunoblotting with the indicated antibodies. Note that RalaG20V does not induce activation of Akt or MAPK. **C and D.** RNA was extracted from S2 cells treated with dsRNA and the levels of indicated targets were measured by qRT-PCR. Graphs show mean normalized to the control (RLuc) +/- stdev from 3 independent experiments.
