## Supplementary figure 6 for "Rala and the exocyst control Pvr trafficking and signaling to ensure lymph gland homeostasis in *Drosophila melanogaster*"

Knævelsrud- Supplementary Figure 6

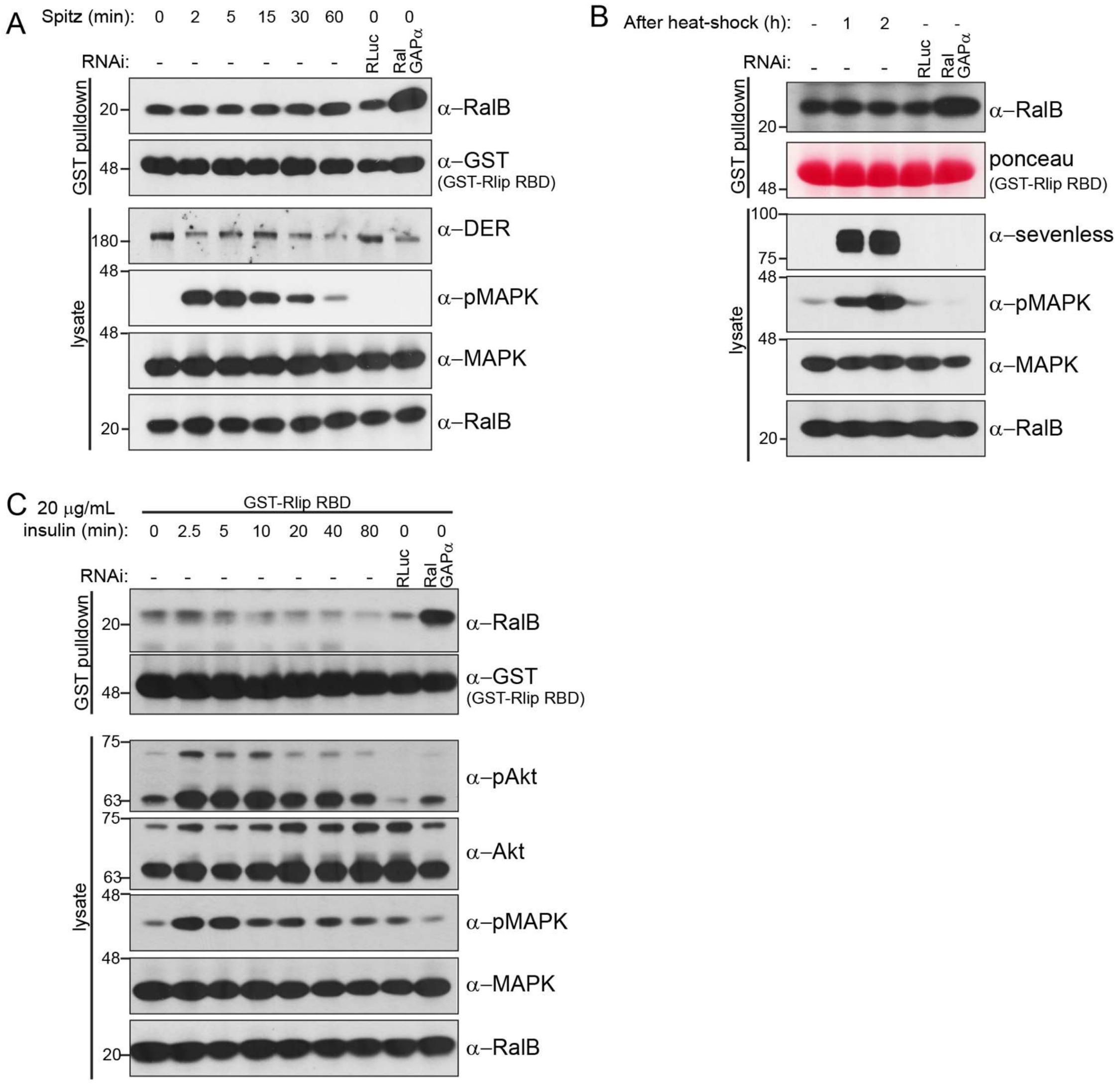


**Supplementary Figure 6. Rala is not activated by active Ras in S2 cells.**

**A.** S2 cells stably transfected with *Drosophila* EGFR (DER) were stimulated for 0 to 60 min with supernatant from Spitz-producing S2 cells. **B.** S2 cells stably transfected with sevenless-S11 under the control of a heat-shock promoter were heat-shocked at 37 °C for 30 min and harvested after 1 or 2 h. **C.** S2 cells were stimulated by 20 µg/mL human recombinant insulin for 0 to 80 min. **A-C.** Recombinant GST-Rlip-RBD bound to GSH sepharose was incubated with the corresponding lysates and the resulting pull-downs analyzed by immunoblotting performed with the indicated antibodies. Activation of the Ras-MAPK pathway was confirmed by increased phospho-MAPK (pMAPK) levels and activation of PI3K-Akt was confirmed by increased phospho-Akt (pAkt) levels.
