## Supplementary figure 7 for "Rala and the exocyst control Pvr trafficking and signaling to ensure lymph gland homeostasis in *Drosophila melanogaster*"

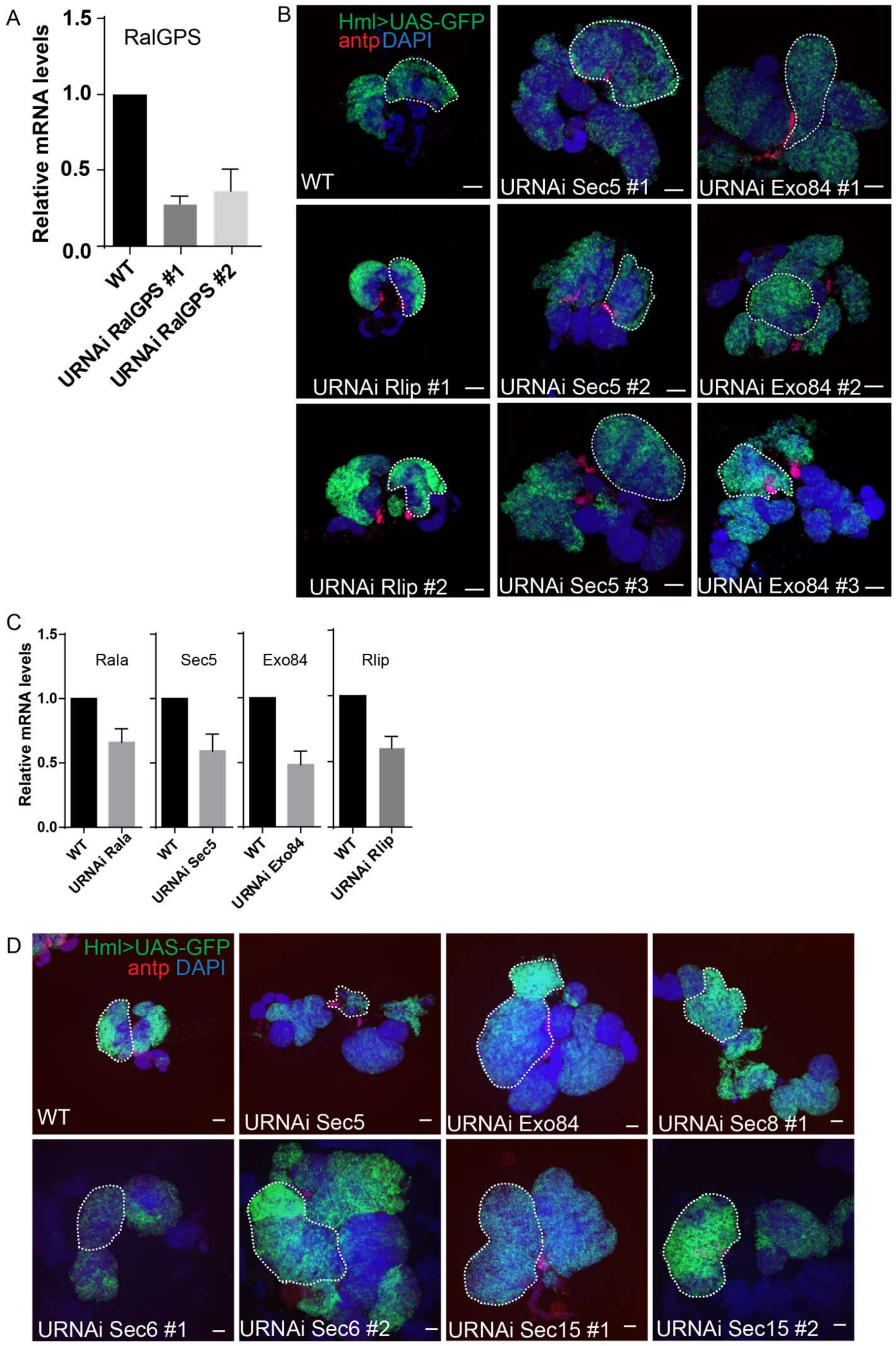


**Supplementary Figure 7. Depletion of the exocyst complex results in enlargement of the lymph gland.**

**A.** RalGPS mRNA levels were measured from entire lymph glands by qRT-PCR. Graphs show mean normalized to the control (WT) +/- stdev from 3 independent experiments. **B and D.** Representative widefield fluorescence images of lymph glands where the cortical zone is marked by Hml-driven GFP and the PSC by antp immunofluorescence. Components of the exocyst complex (Sec5, Exo84, Sec6 or Sec8) or Rlip were depleted by multiple independent RNAi lines. Note that lymph glands where Sec5 is depleted in the cortical zone are sometimes dispersed (D). One primary lobe is outlined in each image. Scale bar 50 µm. **C.** Rala, Sec5, Exo84 and Rlip mRNA levels were measured from entire lymph glands by qRT-PCR. Graphs show mean normalized to the control (WT) +/- stdev from 3 independent experiments.
