## Supplementary figure 8 for "Rala and the exocyst control Pvr trafficking and signaling to ensure lymph gland homeostasis in *Drosophila melanogaster*"

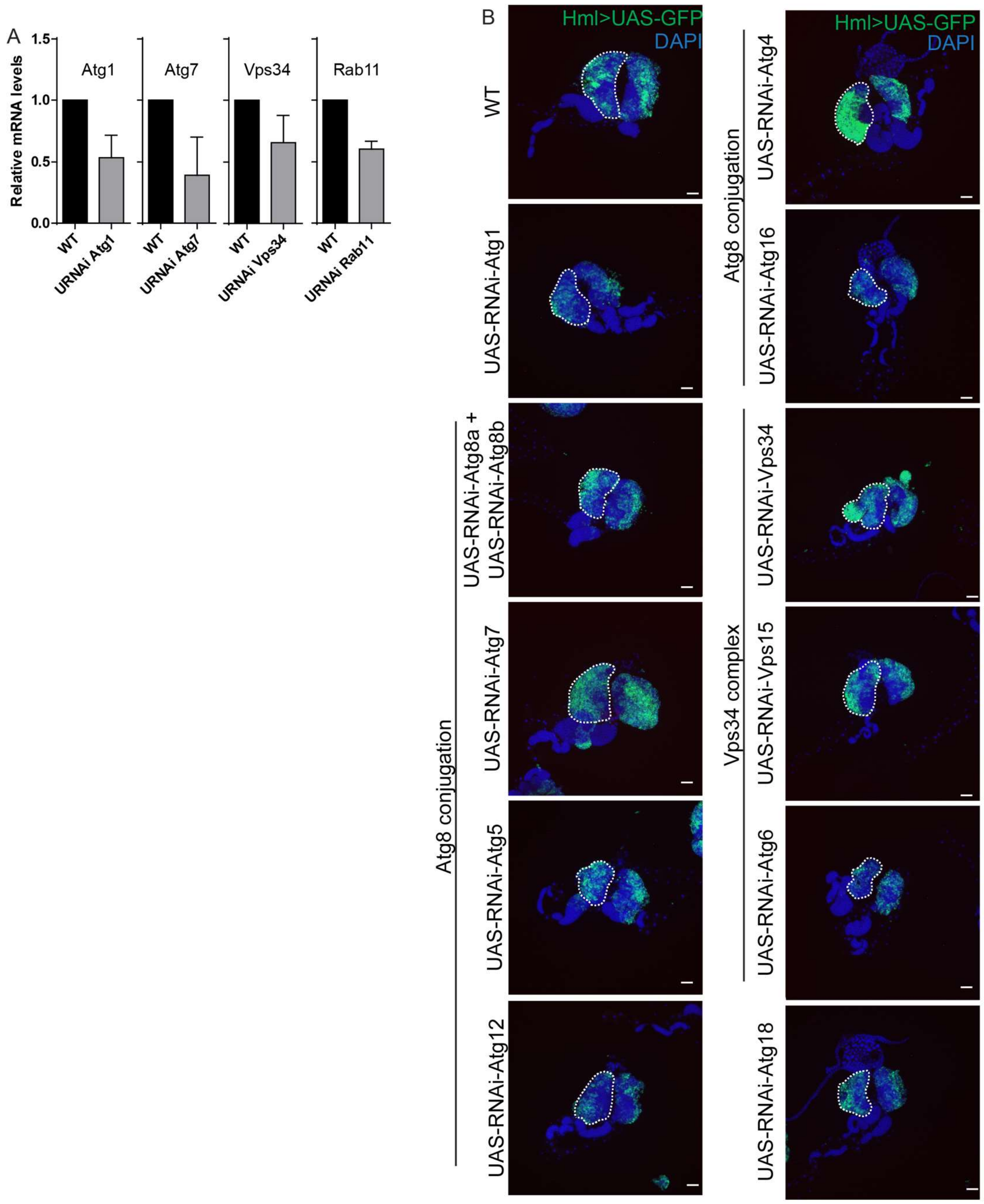


**Supplementary Figure 8. Depletion of autophagy components does not affect lymph gland size.**

**A.** Atg1, Atg7, Vps34 and Rab11 mRNA levels were measured from entire lymph glands by qRT-PCR. Graphs show mean normalized to the control (WT) +/- stdev from 3 independent experiments. **B.** Representative widefield fluorescence images of lymph glands where the cortical zone is marked by Hml-driven GFP and depleted of different components of the autophagy machinery. One primary lobe is outlined in each image. Scale bar 50 µm.
