## Supplementary figure 9 for "Rala and the exocyst control Pvr trafficking and signaling to ensure lymph gland homeostasis in *Drosophila melanogaster*"

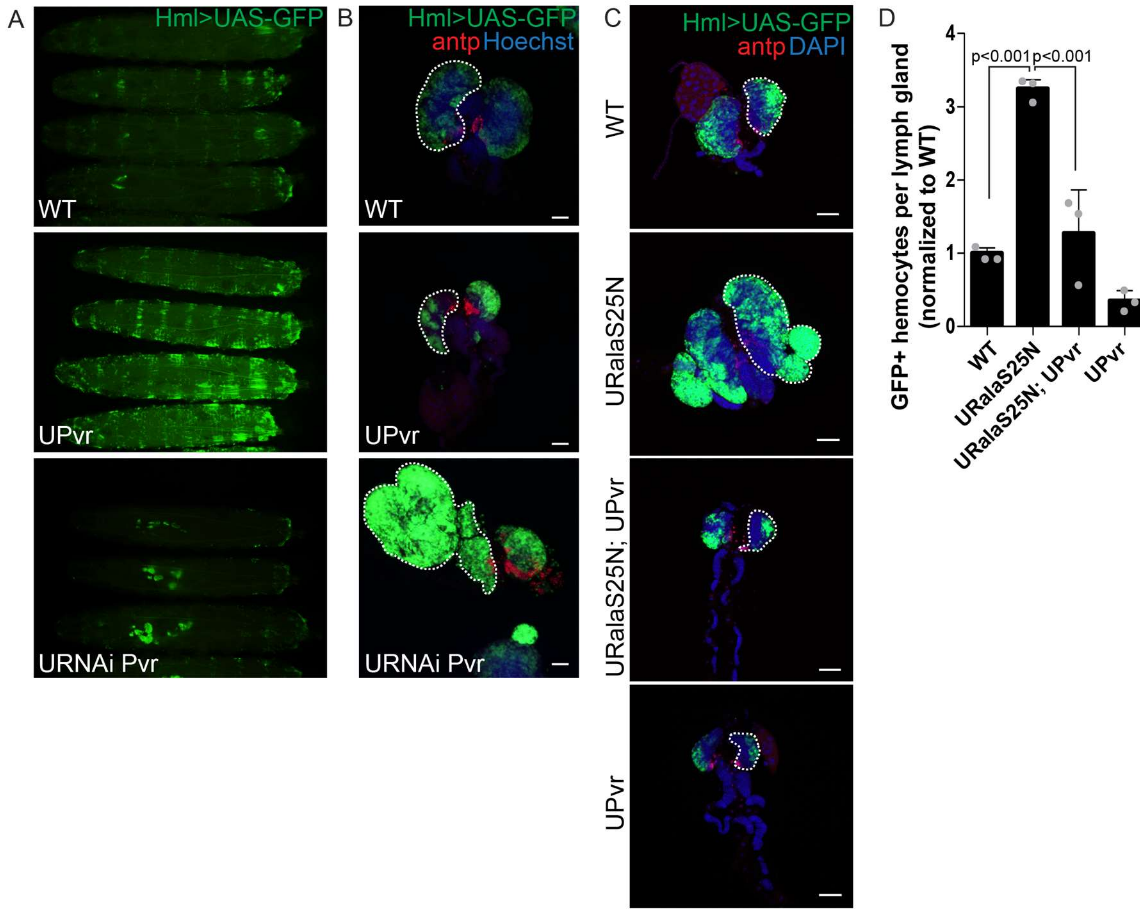


**Supplementary Figure 9. Expression of Pvr rescues RalaS25-induced lymph gland enlargement.**

**A.** Wandering third instar larvae expressing Pvr or dsRNA against Pvr driven by Hml-Gal4 were imaged by widefield fluorescence microscopy to assess the hematopoietic system marked by GFP expression. **B-C.** Representative widefield fluorescence images of lymph glands expressing Pvr or dsRNA against Pvr (B) or dominant-negative Rala alone or in combination with Pvr (C) driven by Hml-Gal4. The cortical zone is marked by Hml-driven GFP and the PSC by antp immunofluorescence. One primary lobe is outlined in each image. Scale bar 50 µm. **D.** Quantification of the number of GFP positive hemocytes per larval lymph gland. Graph shows average GFP-positive hemocytes per lymph gland normalized to control (WT) +/- stdev from 3 independent experiments. Statistically significant differences are indicated (one-way ANOVA).
